# Mechano-dependent sorbitol accumulation supports biomolecular condensate

**DOI:** 10.1101/2023.07.24.550444

**Authors:** Stephanie Torrino, William M Oldham, Andrés R Tejedor, Ignacio S. Burgos, Nesrine Rachedi, Kéren Fraissard, Caroline Chauvet, Chaima Sbai, Brendan P. O’Hara, Sophie Abélanet, Frederic Brau, Stephan Clavel, Rosana Collepardo-Guevara, Jorge R. Espinosa, Issam Ben-Sahra, Thomas Bertero

## Abstract

Biomolecular condensates regulate a wide range of cellular functions from signaling to RNA metabolism^1, 2^, yet, the physiologic conditions regulating their formation remain largely unexplored. Biomolecular condensate assembly is tightly regulated by the intracellular environment. Changes in the chemical or physical conditions inside cells can stimulate or inhibit condensate formation^3–5^. However, whether and how the external environment of cells can also regulate biomolecular condensation remain poorly understood. Increasing our understanding of these mechanisms is paramount as failure to control condensate formation and dynamics can lead to many diseases^6, 7^. Here, we provide evidence that matrix stiffening promotes biomolecular condensation *in vivo*. We demonstrate that the extracellular matrix links mechanical cues with the control of glucose metabolism to sorbitol. In turn, sorbitol acts as a natural crowding agent to promote biomolecular condensation. Using *in silico* simulations and *in vitro* assays, we establish that variations in the physiological range of sorbitol, but not glucose, concentrations, are sufficient to regulate biomolecular condensates. Accordingly, pharmacologic and genetic manipulation of intracellular sorbitol concentration modulates biomolecular condensates in breast cancer – a mechano-dependent disease. We propose that sorbitol is a mechanosensitive metabolite enabling protein condensation to control mechano-regulated cellular functions. Altogether, we uncover molecular driving forces underlying protein phase transition and provide critical insights to understand the biological function and dysfunction of protein phase separation.

## Main

Biomolecular condensates are membraneless assemblies that concentrate biomolecules in cells. Condensed droplets of protein and nucleic acid regulate many cellular functions from signaling to RNA metabolism^1, 2, 6^. The diversity of roles that condensates seem to fulfill is consistent with their stability and physical properties being tightly regulated by the physicochemical parameters (e.g. temperature, pH) and the composition (e.g. concentration of biomolecules, ions, and metabolites) of the intracellular environment^3–5, 8^. However, whether and how the external environment of cells can also regulate biomolecular condensate formation remain unknown.

External mechanical forces control cellular functions ranging from signaling ^9^ to gene regulation ^10^, cell metabolism ^11–15^, and cell proliferation. Mechanotransduction, the process that converts external mechanical signals into intracellular biochemical signals, enables cells to sense and adapt to external mechanical forces. To balance these external forces, cells adjust the stiffness of their cytoskeleton ^9^. This response is achieved through coordinated cytoskeletal rearrangement, actomyosin contraction, and transcription factor activation ^16^, such as the transcriptional cofactors YAP and TAZ (also known as WWTR1). The resulting biomechanical stimuli induce complex and incompletely understood adaptations that influence cellular functions. Therefore, we sought to determine whether and how the mechanical environment of cells can fine-tune biomolecular condensate formation to regulate mechano-dependent cellular functions such as transcription and proliferation.

Here, combining insights from *in silico* multiscale simulations of biomolecular condensates with *in vitro* and *in vivo* phase separation assays, we identified sorbitol as a mechanosensitive metabolite that regulates biomolecular condensate and controls cell proliferation. This is relevant in diseased tissues such as in breast cancer, where increased tissue stiffness is associated with increased sorbitol concentration, increased biomolecular condensates emergence, and increased cell proliferation.

### Matrix stiffening promotes biomolecular condensates phase separations

To determine whether mechanical forces conveyed by the extracellular matrix (ECM) promote protein condensate formation, we investigated the behavior of several proteins known to undergo phase separation^17–19:^ WW domain containing transcription regulator 1 (WWTR1/TAZ; **Fig.1** and **Extended Data Fig.3a**), fused in sarcoma (FUS; **Extended Data** **Fig.1** and **Extended Data Fig.3b**) and galectin-3 (LGALS3/Gal3; **Extended Data** **Fig.2** and **Extended Data Fig.3c**). We cultivated the breast cancer cell line (MDA-MB-231) on hydrogels of varying stiffness, mimicking the mechanical environment of normal mammary gland (1 kPa) or mammary tumors (8 kPa and 50 kPa)^12, 20, 21^. An increased number of cells with condensates, increased condensate intensity, and increased number of condensates per cell were observed in stiff (8 kPa and 50 kPa) conditions with both exogenous (GFP constructs; **Fig. 1a, Extended Data Fig.1a** and **Extended Data Fig.2a**) and endogenous (**Extended Data** **Fig.3**) proteins. A spherical shape and recovery from photobleaching are some of the features of a liquid-like phase-separated structure ^2, 22^. Fluorescence recovery after photobleaching (FRAP) experiments (**Fig.1b, Extended Data Fig.1b** and **Extended Data Fig.2b**), circularity assessment (**Fig.1c, Extended Data Fig.1c** and **Extended Data Fig.2c**), and sensitivity to 1,6-hexanediol (**Extended Data** **Fig.3**), which disrupts liquid-like condensates, confirm a liquid-like phase behavior of the condensates that we observed in cells cultivated on stiff matrix.

**Figure 1:**
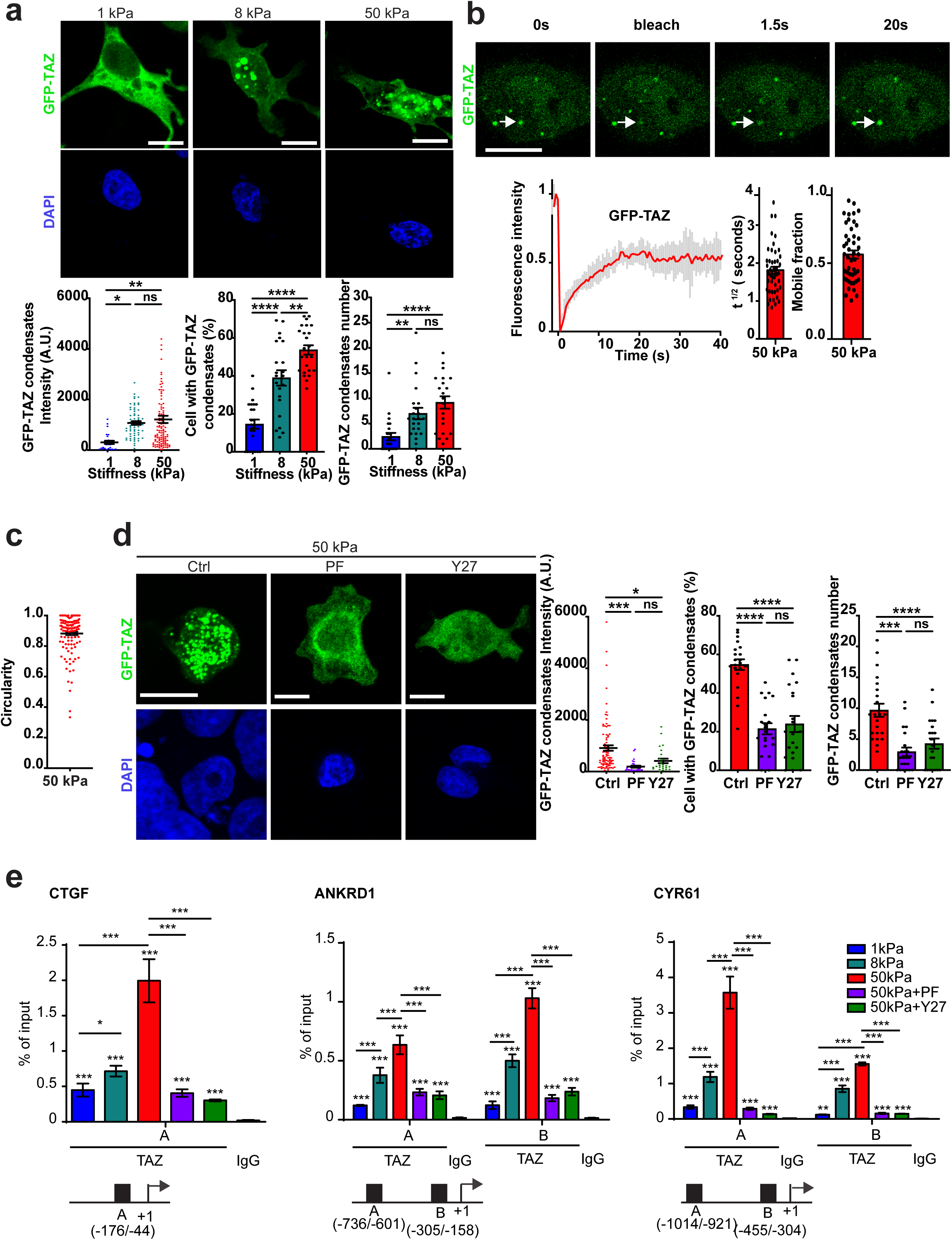
Matrix stiffening promotes TAZ biomolecular condensates. **(a-e)** MDA-MB-231 cells plated on the indicated substrate and treated with 10µM Y-27632 (Y27) or 5µM PF-573228 (PF) for 1 hour **(d-e)**. **(a,d)** Representative immunofluorescence images and quantification of intensity, number per cell of GFP-TAZ condensates and number of cell with condensates. Scale bar=10 µm. **(b)** Representative images and FRAP curves and quantification of diffusion rate (t ½) and mobile fraction of GFP-TAZ. White arrows indicate condensates. **(c)** Circularity quantification of TAZ protein condensates. **(e)** ChIP-qPCR data from cells treated as indicated at CTGF, ANKRD1 and CR61 locus. Results are expressed as percentage of total input DNA prior to immunoprecipitation with anti-TAZ or anti-IgG control. Means of three independent experiments performed in triplicate. In panels a-d n>30 cells from 3 independent experiments were analyzed. ns=not significant; *P<0.05; **P<0.01; ***P<0.001; ****P<0.0001; **(a, d-e)** Bonferroni’s multiple comparison test; data are mean ± s.e.m.

**Figure 2:**
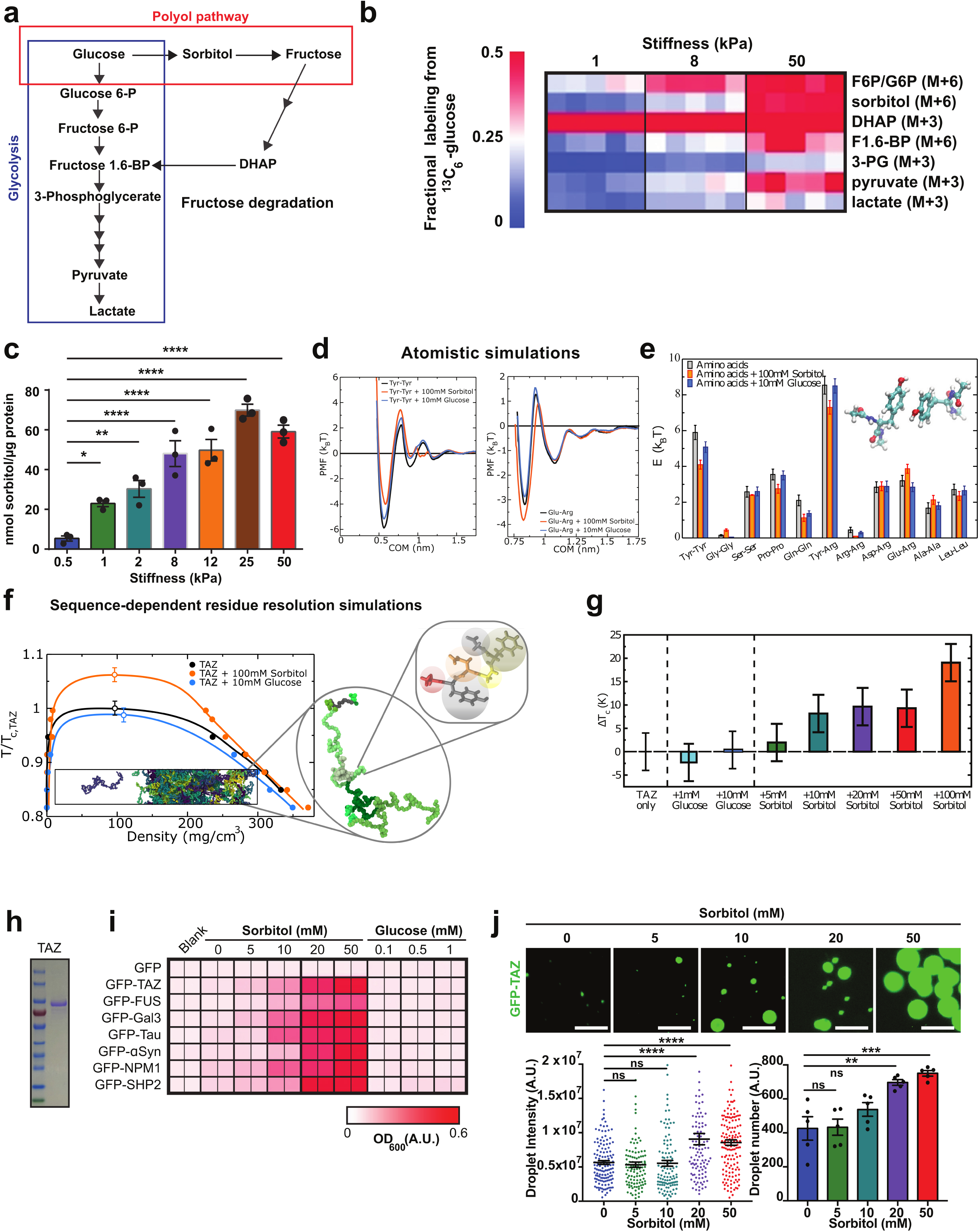
Mechano-induced polyol metabolism regulates biomolecular condensates. **(a)** Schematic representation of glucose pathway**. (b-c)** MDA-MB-231 cells plated on the indicated substrate. **(b)** Heatmap of ^13^C-glucose incorporation in intracellular metabolites. **(c)** Quantification of intracellular sorbitol**. (d)** Atomistic potential of mean force (PMF) simulations for π–π (Tyr–Tyr, left panel) and cation–anion (Glu–Arg, right panel) interactions at physiological NaCl conditions (black curve) and in presence of sorbitol (blue curve) and glucose (orange curve), as a function of the center-of-mass (COM) distance. **(e)** Comparison of the interaction strength of the studied residue-residue pairs from the global minimum in the PMF profiles. **(f)** Phase diagram in the *T* − *ρ* plane for TAZ protein in absence of both sorbitol and glucose (black), in presence of 100 mM of sorbitol (blue), and in presence of 10 mM of glucose using a sequence-dependent residue-resolution model^29^ reparameterized for the simulations with sorbitol and glucose based on the all-atom calculations shown in panel e. Temperatures have been renormalized by the critical temperature of TAZ (312 K) in absence of both solutes. A snapshot of a Direct coexistence simulation of TAZ is included in the phase diagram chart with a zoom-in on a single protein and the coarse-grained representation of each amino acids across the protein sequence. **(g)** Variation of the critical temperature of TAZ from coarse-grained simulations for different concentrations of glucose and sorbitol**. (h)** SDS-PAGE analysis of GFP-TAZ purified protein. **(i)** Heatmap of turbidity analysis of phase separation arising from mixing GFP-TAZ, GFP-FUS, GFP-Gal3, GFP-TAU, GFP-SHP2, GFP-NPM-1, GFP-αSyn or GFP (negative control) with different sorbitol or glucose concentration. **(i)** Representative images and quantification of intensity and number of GFP-TAZ condensates with different sorbitol concentration. Scale bar=10 µm. ns=not significant; **P<0.01; ***P<0.001; ****P<0.0001. **(c,j)** Bonferroni’s multiple comparison test; data are mean ± s.e.m of at least n=3 independent experiments.

**Figure 3:**
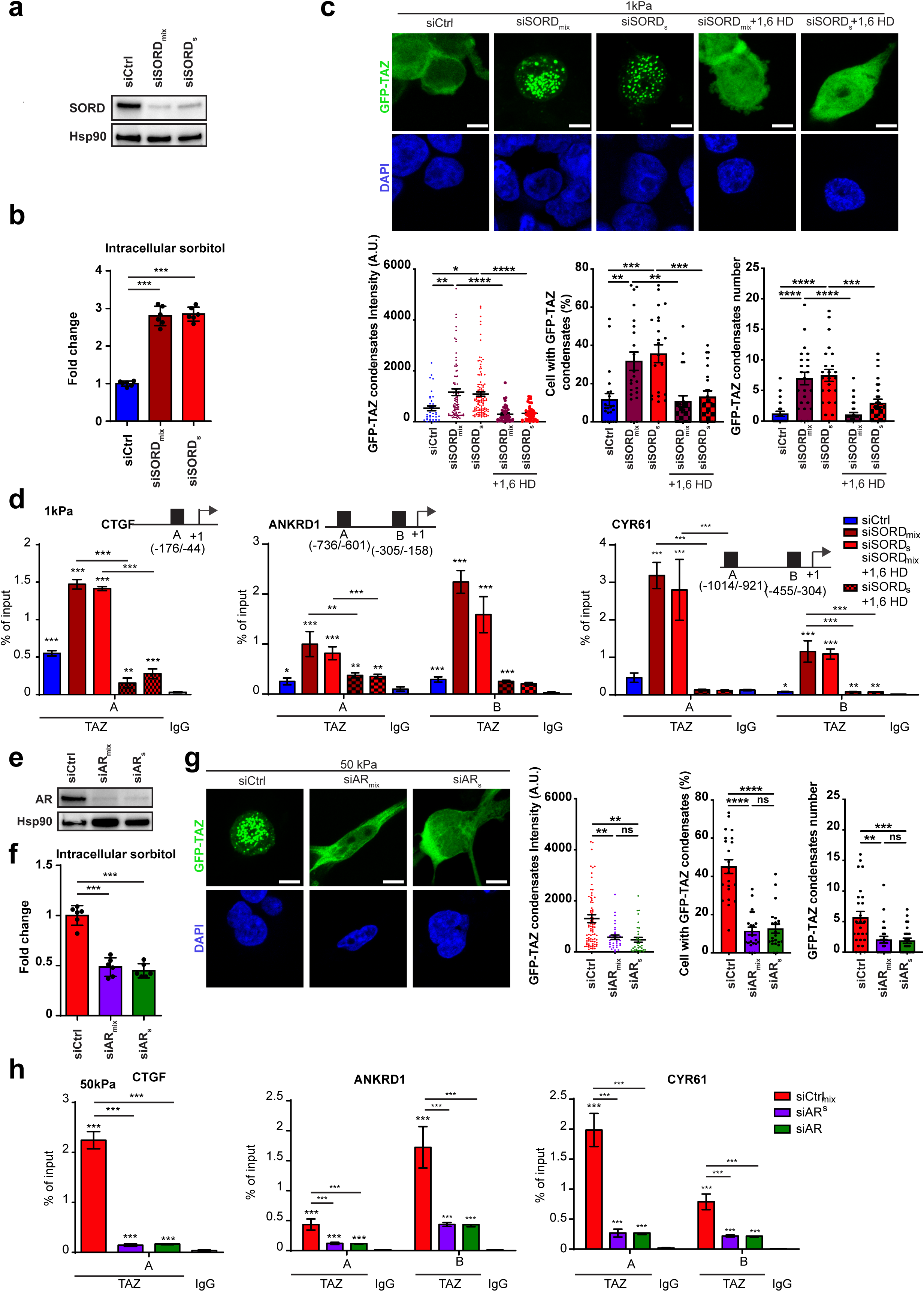
Mechano-dependent biomolecular condensate relies on glucose metabolism to sorbitol. **(a-h)** MDA-MB-231 cells plated on the indicated substrate were transfected with the indicated siRNA (control, siCtrl; SORD and AR, siRNA smarpool or siRNA single) for 48h and treated with 5% of 1,6-Hexanediol (1,6 HD) for 5min **(c-d)**. **(a,e)** Immunoblots of SORD and AR showing the level of siRNA depletion. (**b,f**) Quantification of intracellular sorbitol**. (c,g)** Representative immunofluorescence images and quantification of intensity and number per cell of GFP-TAZ condensates and number of cell with condensates.. In all the panels n>30 cells from 3 independent experiments were analyzed. Scale bar=10 µm**. (d, h)** ChIP-qPCR data from cells treated as indicated at CTGF, ANKRD1 and CR61 locus. Results are expressed as percentage of total input DNA prior to immunoprecipitation with anti-TAZ or anti-IgG control. In all the panels, means of three independent experiments performed in triplicate. ns=not significant; *P<0.05; **P<0.01; ***P<0.001; ****P<0.0001; Bonferroni’s multiple comparison test; data are mean ± s.e.m of at least n=3 independent experiments.

Next, we investigated whether manipulation of the mechanotransduction cascade affects stiffness-dependent biomolecular condensate formation. Cells were cultivated on stiff hydrogels (50 kPa) and treated with either FAK (PF-573228) or Rho-associated kinase (ROCK, Y27632) inhibitors, which attenuate cell contractility similar to growth on soft ECM ^23^. Decreasing cell contractility decreased the number of cells with condensates, condensate intensity, and number of condensates per cell (**Fig.1d, Extended Data Fig.1d** and **Extended Data Fig.2d**). To determine the effect of TAZ condensation on occupancy at target gene promoters ^24^ (CTGF, ANKRD1 and CYR61), chromatin immunoprecipitation (ChIP)-qPCR was performed with antibodies against TAZ in untreated cells cultivated on a gradient of stiffness or cells cultivated on stiff matrix and treated with ROCK or FAK inhibitors. Increasing matrix stiffness increased TAZ at promoters (**Fig.1e)**. Treatments caused a reduction of TAZ at promoters. To demonstrate the broad implication of our findings, we investigated whether mechanical forces could control biomolecular condensate formation in cells from different embryonic origin. Similar results were observed in primary pulmonary arterial endothelial cells (PAECs, **Extended Data** **Fig.4**). Together, our results showed that mechanical cues conveyed by the ECM regulate protein condensate formation.

**Figure 4:**
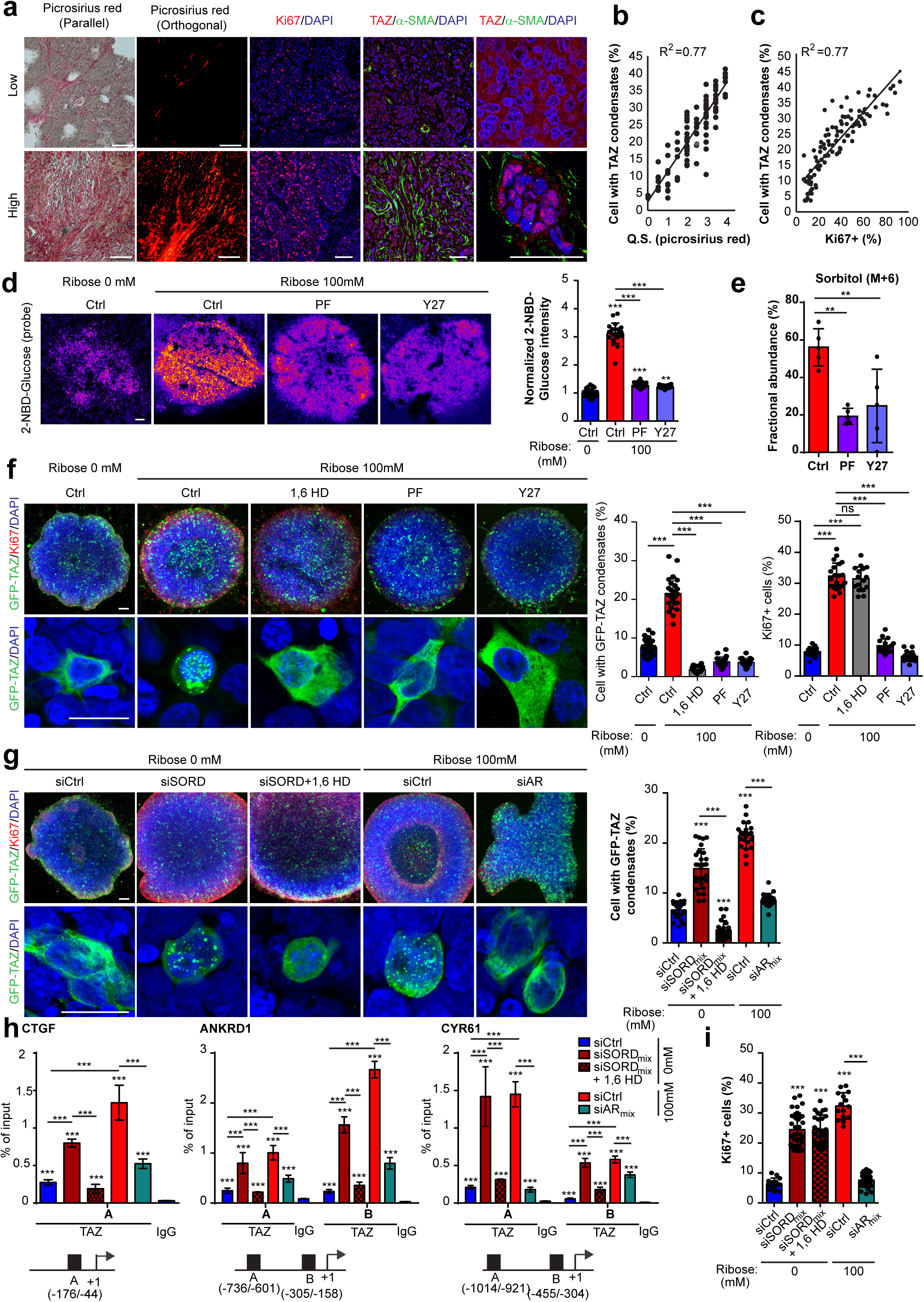
Mechanosensitive polyol pathway regulates biomolecular condensates in breast cancer. **(a)** Representative images of primary mammary tumors are shown as isolated from a cohort of 100 patients. Picrosirius red staining visualized by both parallel and orthogonal light displays tumor ECM remodeling, and representative IHC images show Ki67, α-SMA, and TAZ staining. Scale bar=100 µm. **(b-c)** Correlations between TAZ staining and ECM remodeling **(b)** and between TAZ staining and proliferation **(c)** are shown. Pearson correlation coefficients (R^2^) are indicated. **(d)** Representative immunofluorescence images and quantification of 2-NBD-Glucose intensity of MDA-MB-231 cells cultivated in 3D after the indicated treatment. Scale bar=100 µm. **(e)** Quantification of ^13^C-glucose incorporation in sorbitol in MDA-MB-231 cells cultivated in 3D after the indicated treatment. **(f-g)** Representative immunofluorescence images and quantification of cell with GFP-TAZ condensates and Ki67 of MDA-MB-231 cells cultivated in 3D after the indicated treatment. Scale bar: Upper panel=100 µm and lower panel=20 µm. **(h)** ChIP-qPCR data from cells treated as indicated at CTGF, ANKRD1 and CR61 locus. Results are expressed as percentage of total input DNA prior to immunoprecipitation with anti-TAZ or anti-IgG control. In all the panels n>12 spheroids from 3 independent experiments were analyzed. ns=not significant; **P<0.01; ***P<0.001; **(d-h)** Bonferroni’s multiple comparison test; data are mean ± s.e.m of at least n=3 independent experiments.

### Matrix stiffening mechanoactivates the polyol pathway

Intriguingly, to precipitate phase separation, crowding agents such as polyethylene glycol (PEG), sucrose, ficoll, dextran, or sorbitol are commonly used. Among these agents, sorbitol is a small metabolite from the polyol pathway that can be produced directly by glucose catabolism. Recently, we and others have shown that matrix stiffening rewires cell metabolism to sustain the metabolic needs of mechanoactivated cells^11–14, 20, 25^. Therefore, we asked whether mechanical cues can regulate biomolecular condensate formation through the control of cellular sorbitol levels derived from glucose catabolism. Steady-state metabolomic profiles of cells indicated that ECM stiffening (**Extended Data Fig.5a**) promotes substantial global alterations in cellular metabolism. Pathway analysis revealed that ECM stiffening significantly modulated the steady-state levels of 49 metabolites that are highly enriched in pathways related to glucose metabolism (fructose degradation, glycolysis) (**Extended Data Fig.5b**). Consistent with mechano-induced glycolysis, we observed a dose-dependent correlation between increased matrix stiffness, higher lactate/pyruvate ratio, increased lactate secretion, and elevated glucose consumption (**Extended Data Fig.5c-g**). Moreover, the enhanced glucose uptake coincided with increased expression of the glucose transporter (GLUT1) (**Extended Data Fig.5f)**.

To determine whether stiff matrix exposure redirects glucose to the polyol pathway (**Fig.2a**) we employed stable isotope tracing using [U-^13^C]-glucose where glucose-labelled cells were exposed to matrices with varying stiffness (**Fig.2b** and **Extended Data Fig.5h-m**). Cells cultured on stiff matrices exhibited a higher relative flux through glycolysis compared to those on softer substrates, confirming their increased dependency to glucose consumption. Notably, our results indicated that the glucose-derived sorbitol (M+6) levels increase in response to matrix stiffening (**Fig.2b** and **Extended Data Fig.5i**). Furthermore, we observed a dose-dependent correlation between the degree of matrix stiffness and the intracellular sorbitol concentration (**Fig.2c**). Taken together, these results demonstrate that matrix stiffening enhances glucose consumption, directing it towards the polyol pathway, and leading to an increase in intracellular sorbitol through the activation of the canonical mechanotransduction pathway.

### Multiscale simulations predict that sorbitol—but not glucose**–**promote biomolecular condensate formation

To further understand the molecular mechanism explaining the effect of sorbitol and glucose in biomolecular condensates stability, we developed a multiscale modelling approach that leverages molecular dynamics (MD) simulations at two complementary resolutions. Our highest resolution corresponds to atomistic potential-of-mean-force (PMF) simulations^26^; these level of modelling allows us to quantify the relative strength of amino acid pair interactions at different sorbitol and glucose concentrations representative of the cell environment **(Fig.2d-e and Extended Data Fig.6**; simulation details are provided in the **Methods Section**). We then decrease the resolution of the models to reach the spatiotemporal scales needed to investigate condensate behaviour by mapping our atomistic results into a residue-resolution coarse-grained model. Such residue-resolution model allows us to perform simulations of TAZ protein condensates and compare condensate stability in the presence and absence of sorbitol (**Fig. 2f-g**).

Our atomistic simulations reveal that the effective interaction strength between aromatic residues, i.e. Tyrosine–Tyrosine, and among polar residues, i.e. Glutamine–Glutamine, are weaker in the presence of glucose (10 mM) than in its absence. The strengths of other associative interactions (e.g., cation–π, charge–charge, and hydrophobic) are negligibly affected by low concentrations of glucose (i.e., ≤ 10 mM; **Fig.2e**). Such modulation of amino acid pair interactions by glucose is consistent with inhibition of phase separation, mainly because of π – π interactions being chief drivers of biomolecular phase separation of intracellular proteins at physiological conditions^27, 28^. We then turn our attention to sorbitol, which is present at significantly higher concentrations than glucose upon matrix stiffening through the polyol pathway. Our simulations reveal the striking molecular mechanism by which sorbitol enhances the formation of biomolecular condensates. Sorbitol asymmetrically transforms the quality of the solvent: (a) increasing the affinity of the solvent for π-rich residues, and hence weakening π–π and cation–π bonds, and (b) decreasing the affinity of the solvent for charged amino acids and spacers, hence enhancing the strength of associative electrostatic interactions and spacer–spacer interactions. Specifically, we observe that the effective interaction strength among Tyrosine–Tyrosine, Tyrosine–Arginine, and Proline– Proline decreases in the presence of 100 mM sorbitol **(Extended Data Fig.6**). In contrast, the strength of electrostatic interactions (such as arginine-glutamic acid or arginine-aspartic acid) as well as between highly abundant weak-interacting amino acids within proteins (e.g., glycine or alanine) moderately increase in presence of 100 mM of sorbitol (**Fig.2e**).

To macroscopically elucidate the overall impact of both sorbitol and glucose in TAZ condensate stability, we map our all-atom PMF binding energies (**Fig.2e**) into a residue-resolution coarse-grained (CG) model that accurately describes the experimentally observed conformational propensities and phase-separation behavior of a large set of intrinsically disordered proteins at physiological conditions^29^ (further details on the CG implementation and the sorbitol/glucose reparameterization of the model are provided in the Methods). By performing Direct Coexistence (DC) simulations with this model^30^, we evaluate the phase diagram in the temperature-density plane of TAZ (and FUS; **Extended Data Fig.7a**) at physiological salt concentration but in the absence of glucose and sorbitol (black curve; **Fig.2f**), and compare with the results of adding low glucose concentrations (blue curve) vs. high sorbitol concentration (orange curve). In our CG simulations, we consider hundreds of protein replicas. Importantly, we detect that the critical temperature of TAZ—which is a direct measurement of condensate stability^31^—is enhanced by the addition of sorbitol but not glucose **(Fig.2f)**. The same behavior is found for FUS condensates where glucose (at concentrations lower than 10 mM) barely changes condensate stability, whereas sorbitol significantly augments it (**Extended Data Fig.7a)**. By analysing the influence of different concentrations of glucose and sorbitol in TAZ condensate stability through simulations, we conclude that sorbitol is only able to enhance protein condensate stability once its concentration is above 10 mM (**Fig.2g**).

### Physiological levels of sorbitol promote biomolecular condensates *in vitro*

Based on our multiscale simulations **(Fig.2d-g** and **Extended Data Fig.6-7**), we wanted to determine whether the physiological increase in sorbitol indeed promoted protein condensates (**Fig.2h-j and Extended Data Fig.8**). To test this idea, we performed an *in vitro* phase separation assay of a wide range of proteins, including TAZ, FUS, and Gal3, as well as Tau, α-synuclein (α-Syn), Nucleophosmin 1 (NPM1), and SH2-containing protein tyrosine phosphatase 2 (SHP2), all known to undergo phase separation^32–35^. We purified GFP-tagged proteins (**Fig.2h and Extended Data Fig.8a**) and added them to sorbitol across a range of physiological concentrations (as determined in **Fig.2c**) and assessed whether these proteins form condensates *in vitro*. Consistent with *in silico* models, we found that increasing sorbitol, but not glucose, increased the solution turbidity of all these proteins. By contrast, no increase in turbidity was observed for purified GFP control (**Fig. 2i**). Furthermore, as visualized by confocal imaging, purified GFP-tagged proteins spontaneously formed micro-sized droplets in solution (**Fig. 2j and Extended Data Fig.8b-g)**, and the droplets were more numerous with greater intensity at higher sorbitol concentrations. Together, these results in combination with our simulations, demonstrate that increasing sorbitol across physiological concentrations is sufficient to promote protein condensates.

### Mechano-dependent biomolecular condensates rely on intracellular sorbitol

To test whether mechano-induced protein condensates depend on the mechanosensitive polyol pathway *in vivo*, we performed a series of complementary experiments in which we manipulated the intracellular sorbitol concentration in cells cultivated on soft or stiff substrates. We cultivated MDA-MB-231 cells on soft substrate while silencing sorbitol dehydrogenase (SORD; siSORD) expression. SORD converts sorbitol to fructose and its inhibition increases intracellular sorbitol. Consistent with our hypothesis, siRNA knockdown of SORD (**Fig.3a**) in cells cultivated on soft substrate increases intracellular sorbitol **(Fig.3b)** and promotes GFP-TAZ (**Fig.3c**) protein condensates. Importantly, treatment with 1,6-hexanediol reduced the number of TAZ condensates. ChIP-qPCR revealed increased TAZ occupancy at target gene promoters in SORD knockdown cells, which was decreased in 1,6-hexanediol-treated cells (**Fig.3d**), highlighting the functional relevance of these findings. Conversely, we investigated whether pharmacologic (Ethyl 1-benzyl-3-hydroxy-2(5H)-oxopyrrole-4-carboxylate (EBPC), **Extended Data Fig.9**) or genetic inhibition of aldose reductase (AR; siAR, **Fig.3e**), the enzyme that converts glucose to sorbitol, decreased intracellular sorbitol (**Fig.3f**) and impacted protein condensates (**Fig.3g**). Inhibition of AR in cells cultivated on stiff substrate decreased GFP-TAZ (**Fig. 3g and Extended Data Fig.9a**), GFP-FUS (**Extended Data Fig.9b**) and GFP-Gal3 (**Extended Data Fig.9c**) protein condensates. We also observed these effects in PAECs (**Extended Data Fig.9d-f**). Consistent with these results, ChIP-qPCR reveals decreased TAZ occupancy at the promoter of target genes in AR knockdown cells (**Fig.3h**). Thus, manipulating the polyol pathway fine-tunes proteins condensate formation *in vivo*.

### Matrix stiffening promotes sorbitol-dependent protein condensates in breast cancer models

We next sought to establish the pathophysiologic relevance of our findings (**Fig.4** and **Extended Data Fig.10**). Given the role of breast tumor niche remodeling and stiffening in both cancer cell metabolism ^12^ and tumor progression ^36, 37^, we investigated whether protein condensates are observe in a cohort of 100 human patients with a pathological diagnosis of breast cancer ^20^. As tumor niche stiffening is known to correlate with collagen deposition and assembly in breast cancer ^36^, we quantified collagen deposition and assembly by picrosirius red staining. In remodeled (picrosirius red positive) and proliferative (Ki67+ cells) tumors, a concurrent upregulation of protein condensates was observed (**Fig.4a-c** and **Extended Data Fig.10a-b**), indicating that protein condensates are associated with proliferative and remodeled breast cancer tumors. Based on these findings, we investigated whether tumor niche stiffening modulates tumor cell metabolism to promote protein condensates in an orthotopic syngeneic mouse model of breast cancer (**Extended Data Fig.10c-j**). Using a known pharmacologic inhibitor (β-aminopropionitrile, BAPN) of lysyl-oxidase (Lox)^36, 37^, the enzyme responsible for collagen cross-linking and consequent matrix stiffening, we determined whether decreased mechanical cues conveyed by the ECM could prevent metabolic changes and protein condensates. Consistent with our results, decreased matrix stiffness by BAPN (**Extended Data Fig.10d**) decreased intratumoral sorbitol (**Extended Data Fig.10e**). Such metabolic effects further decreased protein condensates (**Extended Data Fig.10f-g, i and j**). Importantly, decreased appearance of TAZ condensates is correlated with lower tumor cell proliferation (**Extended Data Fig.10h**). Together our results indicate that matrix stiffening increased both intratumoral sorbitol and emergence of protein condensates in breast cancer.

Assessing whether mechanosensitive changes in sorbitol promoted protein condensate formation to sustain mechano-dependent cellular activities *in vivo* remains challenging. Therefore, to gain further insight into the molecular mechanisms in a pathophysiologic mechanical context, we performed a series of experiments in a 3D tumoroid model in which we can manipulate the mechanical environment (**Fig.4d-i**). First, we cultivated MDA-MB-231 cells as tumoroids in a low collagen microenvironment (1 mg/mL) and manipulated the mechanotransduction cascade using either ribose (100 mM), a FAK inhibitor, or a ROCK inhibitor (**Fig.4d**). Addition of ribose increases glucose uptake and collagen stiffness^12^. Pharmacologic inhibition of FAK or ROCK decreased glucose uptake (**Fig.4d**). Importantly, decreased glucose uptake following FAK or ROCK inhibition is associated with decreased glucose-derived sorbitol in tumoroids as demonstrated by ^13^C6-glucose tracing (**Fig.4e**). Second, using MDA-MB-231 cells that stably expressed GFP-TAZ, we observed that addition of ribose increased the number of GFP-TAZ cells with condensates, and increased the number of proliferative cells (**Fig.4f**). Conversely, pharmacologic inhibition of FAK or ROCK decreased the number of GFP-TAZ cells with condensates (**Fig. 4f**) and decreased the number of proliferative (Ki67+) cells (**Fig.4f**). Importantly, treating cells with 5% of 1,6-hexanediol reduced the number of cells with TAZ condensates. Third, to investigate whether the mechanosensitive polyol pathway is required to promote TAZ condensates and sustain mechano-dependent cell activities, such as transcription and proliferation, we manipulated the intracellular sorbitol concentration in tumoroids (**Fig.4g-i**). In a low collagen microenvironment, siRNA knockdown of SORD promoted GFP-TAZ protein condensates (**Fig.4g**), increased TAZ occupancy at target genes promoter (**Fig.4h**), and increased cell proliferation (**Fig.4i**). Addition of ribose to increase collagen stiffness also increased the number of cells with GFP-TAZ protein condensates (**Fig.4g**), increased TAZ occupancy at target genes promoter (**Fig.4h**), and increased cell proliferation (**Fig.4i)**, features that were blunted by AR inhibition. Importantly, treatment with 1,6-hexanediol caused a reduction in the number of cells with GFP-TAZ condensates (**Fig.4g**) and decreased TAZ occupancy at target genes promoter (**Fig.4h**). Together our results delineate the mechanosensitive sorbitol metabolite as a direct regulator of protein condensates to sustain mechano-induced cell activities in breast cancer models.

In summary, our findings establish mechanical forces as direct modulators of protein condensates. We identified the mechanosensitive polyol pathway as a natural mediator of cytoplasmic and nuclear protein condensates through tightly controlled intracellular sorbitol accumulation. Based on atomistic and coarse-grained simulations, showing that a physiological range in sorbitol, but not glucose, concentrations, is sufficient to regulate biomolecular condensates, we demonstrated that manipulating sorbitol concentration is sufficient to promote protein condensates *in vitro* and i*n vivo*. Finally, we established the pathophysiological relevance of our findings. We demonstrate that the mechanosensitive sorbitol metabolite promotes biomolecular condensate to sustain mechano-induced cell activities in breast tumor—a mechano-dependent disease. Together, these results carry broad implications for our fundamental understanding of both the regulation of biomolecular condensate formation and the mechano-metabo connection in health and disease.

Matrix stiffening fuels aberrant cell behavior and is a hallmark of diseases progression, ranging from cancer, fibrosis, and cardiovascular diseases to neurological defects ^10, 11, 36, 38, 39^. Recently, we and other have shown that ECM stiffening control glucose homeostasis in both transformed ^12, 14^ and non-transformed cells ^11, 40^. Although we have observed increased glucose consumption and lactate fermentation in response to ECM stiffening in highly glycolytic cells such as cancer cell lines and diseased vascular cells, the regulatory pathways allowing this metabolic adaptation have remained enigmatic. Here, we show that matrix stiffening reroutes a large amount of glucose to the polyol pathway (**Fig.2**). Therefore, it is tempting to speculate that by bypassing hexokinase, the first and a rate-limiting reaction in glycolysis ^41^, the conversion of glucose to fructose through the polyol pathway allows highly glycolytic cells to promote fructolysis, and thus sustain their important metabolic demands. As such, matrix stiffening not only promotes glycolysis, but also promotes fructose degradation making proliferative cells such as cancer cells even more proliferative (**Fig.4**). Intriguingly, sorbitol and its derivatives have also been used by the food industry as artificial sweeteners in a wide range of foods and beverages. The safety of artificial sweeteners remains debated, with conflicting findings regarding their role in carcinogenesis ^42^. Therefore, and as suggested by our study, whether these artificial sweeteners modulate protein phase behavior has to be determined. Further studies should also determine whether chronic exposure to these artificial sweeteners modulates protein phase behavior in humans.

We also have shown that mechanoregulated glucose catabolism through the polyol pathway increases intracellular sorbitol. Sorbitol has been used as a crowding agent. Here, we demonstrated that variation of sorbitol concentration in a physiologic range is sufficient to modulate protein condensates. Although the concentration of other small metabolites, such as ATP, has been proposed to regulate biomolecular condensates^5^, their physiological relevance has remained poorly studied. Here, we show that sorbitol is directly produced from glucose catabolism and thus rapidly mobilized to promote endogenous protein condensates. Intracellular sorbitol concentration is tightly controlled by mechanical stress. Indeed, our glucose tracing experiments reveal that the intracellular pool of sorbitol derived from glucose remains stable over time and that increasing matrix stiffness gradually increased intracellular sorbitol concentration (**Fig.2**). Furthermore, pharmacologic manipulation of the polyol pathway reveals that sorbitol-dependent protein condensation is highly dynamic. Inhibition of sorbitol production in cells cultivated on stiff matrix using EBPC decreased proteins condensates in less than thirty minutes. Together, these results place sorbitol as the ideal metabolite to regulate biomolecular condensate in (patho)physiological conditions. Further studies are necessary to determine whether other environmental stresses can also impact polyol metabolism and consequent biomolecular condensates.

Finally, biomolecular condensates have been implicated in diverse biological processes, including chromatin reorganization^43^, noise buffering^44^, and sensing^45^, and their misregulation has been associated with the emergence of diverse pathologies, such as neurodegenerative diseases and cancer^6, 7, 17^. Here we highlight a physiological mechanism regulating biomolecular condensate formation. As such, we not only uncover molecular driving forces underlying protein phase transitions, but also provide critical insight to understand the biological function and dysfunction of protein phase separation. The polyol pathway and matrix stiffening are dysregulated in many diseases including cancer and cardiovascular diseases. Accordingly, we provide fundamental insight regarding the role of biomolecular condensates in breast cancer. We found that biomolecular condensates are associated with tumor cell proliferation and increased ECM remodeling in a cohort of breast cancer patients. Consistent with the role of TAZ in self-renewal and tumor-initiation in breast cancer^46^ and the role of TAZ condensate in the transcriptional activation^18^, we demonstrate that mechano-induced TAZ condensate sustains TAZ transcriptional activity and subsequent cell proliferation in models of breast cancer. Therefore, our results open new avenues to investigate the role of the mechanosensitive polyol pathway and its influence in biomolecular condensate related-diseases.

## Online Methods

### Cell culture

MDA-MB-231 cells were purchased from the American Type Culture Collection (ATCC). Cells used in this study were within 20 passages after thawing and were cultured (37°C, 5% CO2) in Dulbecco’s Modified Eagle’s Medium (DMEM, Gibco) supplemented with 10% fetal bovine serum (Gibco), Glutamine (2mM, Gibco) and Penicillin/Streptomycin (1%, Gibco). Primary human (Lonza) pulmonary arterial endothelial cells (PAECs) were grown in EBM-2 basal medium supplemented with EGM-2 MV BulletKit (Lonza). For the studies dependent on matrix stiffness, collagen-coated hydrogel pre-plated in culture wells (Matrigen) was generated from a mix of acrylamide and bis-acrylamide coated with collagen. Cells were cultured, passaged, and harvested while on top of the hydrogel, using standard cell culture techniques.

### Plasmids, antibodies and reagents

pEGFP-FUS (#60362), pEGFP-TAZ (#66850), pEGFP-TAU (#46904), pEGFP-α-Syn (#40822), pEGFP-NPM1 (#17578), pEGFP-SHP2 (#12283) and pEGFP-Gal3 (#73080) plasmids were purchased from Addgene. Theses plasmids were double-digested and ligated with pET28a (+) (EMDMillipore). Constructs were verified by sequencing.

D-Sorbitol (S7547), D-Glucose (G8270), D-Ribose (R9629), Y-27632 (SCM075), PF-573228 (PZ0117), EBPC (PZ0140) and 1,6-Hexanediol (240117) were purchased from Sigma. DTT (10699530) was purchased from Thermo Fisher Scientific. The following commercially available antibodies were used for western blotting and immunofluorescence: - mouse monoclonal antibodies against Galectin-3 (Proteintech, 60207-1-Ig), Aldose reductase (Santa cruz biotechnology, SC-271007), GFP (Roche, 11814460001) and Hsp90 (Santa Cruz Biotechnology, sc-69703); - rabbit polyclonal antibodies against FUS (Abcam, ab23439), TAZ (Santa-Cruz, sc-48805), Ki67 (Abcam, ab15580) and SORD (Altas antibodies, HPA040621). HRP-conjugated donkey anti-mouse IgG (715-035-150) and HRP-conjugated anti-mouse IgG (711-035-152) were purchased from Jackson ImmunoResearch Laboratories.

### Protein expression and purification

pET28a-EGFP-FUS, pET28a-EGFP-TAZ, pET28a-EGFP-TAU, pET28a-EGFP-α-Syn, pET28a-EGFP-NPM1, pET28a-EGFPSHP2 and pET28a-EGFP-Gal3 were transformed into *E.coli* BL21 (Invitrogen) cells using standard protocol. Cells were grown to OD 0.6, lyzed and purified using MagBeads Ni-IMAC Pierce high-capacity (Thermo Fisher Scientific) according to the manufacturer’s instructions. The eluted proteins were dialyzed overnight at 4 °C and concentrated with Amicon Ultra Centrifugal Filters (Millipore). Clean proteins were verified with SDS PAGE.

### Turbidity assay

Purified protein was mixed with various sorbitol or glucose concentrations in a 50 μL total volume, containing 20mM Tris-HCl pH7.5, 50mM NaCl and 1mM DTT, in a Greiner 96 well clear microplate. The protein concentration was 5 µM for FUS, 50 µM for TAZ, 40 µM for Gal3, 10 µM for Tau, 8 µM for SHP2, 200 µM for α-Syn, and 20 µM for NPM1. Sample were incubated at room temperature for 10 minutes prior to the absorption (turbidity) measurement at 600nm in a microplate reader. Readings were recorded in duplicate for each protein sample.

### Droplet assay

Purified proteins were subjected to a series of sorbitol concentrations in a total volume of 20 μL containing 20mM Tris-HCl pH7.5, 50mM NaCl and 1mM DTT. Samples were deposited on µ-Slide 18 Well - Flat Ibidi slides (81826), incubated at room temperature for 5 minutes before being imaged on Cytation5, Biotek. ImageJ software was used in all image processing.

### Three-dimensional cell culture

Spheroid assay was performed as we described before^12, 20^. Briefly, MDA-MB-231 cells were removed from the cell culture dishes with trypsin and re-suspended in DMEM 10% FCS at a concentration of 1 × 10^6^ cells per mL. Fifty-microliter droplets were plated onto the underside of a 10 cm culture dish and allowed to form spheroids in a 37 °C incubator for 24 hours. The tumoroids were then embedded in a collagen I/Matrigel gel mix at a concentration of approximately 1 mg/mL collagen I and 2 mg ml^−1^ Matrigel (BD Bioscience) in 24-well glass-bottomed cell culture plates (MatTek). The gel was incubated for at least 45 min at 37 °C with 5% CO2. The gel was covered with DMEM media

### siRNA and Plasmid transfection

Cells were plated on collagen-coated plastic (50μg/mL) and transfected 24h later at 70-80% confluence using siRNA (25nM) or plasmid (3µg/ 10^6^ cells) and Lipofectamine 2000 reagent (Life Technologies), according to the manufacturers’ instructions. Eight hours after transfection, cells were trypsinized and re-plated on hydrogel or used for spheroid assay. siRNA ON-TARGETplus Human SORD (pool: L-004036-00-0005; single: J-004036-05-0005), Human AKR1B1 (pool: L-004036-00-0005; single: J-004036-07-0005) or Non-Targeting Control siRNAs (D-001810-01) were purchased from Horizon Discovery Ltd.

### Lentivirus production

HEK293T cells were transfected using Lipofectamine 2000 (Life Technologies) with lentiviral plasmids along with packaging plasmids (pPACK, System Biosciences), according to the manufacturer’s instructions. Virus was harvested, sterile filtered (0.45-μm), and utilized for subsequent infection of MDA-MB-231 (24-48 hour incubation) for gene transduction.

### Immunoblot assays

Forty-eight hours after plaiting, cells were lysed in RIPA buffer (Pierce) or directly in Laemmli’s buffer. After denaturation, protein lysates were resolved by SDS-PAGE and transferred onto a PVDF membrane (Millipore). Membranes were blocked with 2% BSA in TBS tween20 0.1% and incubated in the presence of the primary and then secondary antibodies. After washing, immunoreactive bands were visualized with ECL (Millipore) and analyzed on Fusion-FX Imager (Vilber).

### Messenger RNA extraction

Cells were homogenized in 1 mL of QiaZol reagent (QIAGEN). Total RNA content was extracted using the miRNeasy kit (QIAGEN) according to the manufacturer’s instructions. Total RNA concentration was determined using a ND-1000 micro-spectrophotometer (NanoDrop Technologies).

### Quantitative RT-PCR of messenger RNAs

Messenger RNAs were reverse transcribed using the Multiscript RT kit (Life Technologies) to generate cDNA. cDNA was amplified via fluorescently labeled Taqman primer sets using an Applied Biosystems 7900HT Fast Real Time PCR device. Fold-change of RNA species was calculated using the formula (2-ΔΔCt), normalized to RPLP0 expression.

### Targeted LC-MS

Metabolite extraction was performed essentially as described^12, 20^ with minor modifications. Briefly, metabolites were extracted from cultured cells on dry ice using 80% aqueous methanol precooled at –80°C. Insoluble material from both cell and supernatant extractions was removed by centrifugation at 20,000 g for 15 minutes at 4°C. The supernatant was evaporated to dryness by SpeedVac at 42 °C, the pellet was resuspended in LC-MS water, and metabolites were analyzed by LC-MS.

LC-MS analysis was performed on a Vanquish ultra-high performance liquid chromatography system coupled to a Q Exactive mass spectrometer (Thermo) that was equipped with an Ion Max source and HESI II probe. External mass calibration was performed every seven days. Metabolites were separated using a ZIC-pHILIC stationary phase (150 mm × 2.1 mm × 3.5 mm; Merck) with guard column. Mobile phase A was 20 mM ammonium carbonate and 0.1% ammonium hydroxide. Mobile phase B was acetonitrile. The injection volume was 1 μL, the mobile phase flow rate was 100 μL/min, the column compartment temperature was set at 25°C, and the autosampler compartment was set at 4 °C. The mobile phase gradient (%B) was 0 min, 80%; 5 min 80%; 30 min, 20%; 31 min, 80%; 42 min, 80%. The column effluent was introduced to the mass spectrometer with the following ionization source settings: sheath gas 40, auxillary gas 15, sweep gas 1, spray voltage +/-3.0 kV, capillary temperature 275 °C, S-lens RF level 40, probe temperature 350 °C. The mass spectrometer was operated in polarity switching full scan mode from 70-1000 m/z. Resolution was set to 70,000 and the AGC target was 1×10^6^ ions. Data were acquired and analysed using TraceFinder software (Thermo) with peak identifications based on an in-house library of authentic metabolite standards previously analysed utilizing this method. For all metabolomic experiments, the quantity of the metabolite fraction analysed was adjusted to the corresponding cell number calculated upon processing a parallel experiment.

### Stable isotopic tracing analysis

For collecting cells from 10-cm plates, cells were washed with medium 5 ml with glucose-free medium and incubated with the tracer (^13^C_6_-glucose, 20 mM) for 1 hour or 24 hours prior to metabolite collection. All media were removed and plates were kept tilted for few seconds to allow any additional media collection in corner of plate. Cells were immediately treated with 4 ml of 80% ice-cold methanol (-80°C) and transferred to −80°C freezer for 20 minutes. Cell plates were resuspended on dry ice and the collected lysate/methanol suspension was transferred to 15 ml conical tubes kept on dry ice. Suspension was centrifuged at full speed (21,000 *g*) for 5 minutes at 4°C. Supernatants were then transferred to 50 ml conical tubes on dry ice. The pellets were resuspended with 500 μl 80% methanol (-80°C) and a combination of vortexing and pipetting up and down. The suspension was centrifuged at full speed (21,000 *g*) for 5 minutes at 4°C. Supernatant were transferred to 50 ml conical tubes on dry ice. Pellets obtained after centrifugation were dissolved in urea 8 M (Tris 10 mM pH 8.0) at 60°C, to later measure protein concentration for normalization. After pooling the metabolite extractions, the samples are completely dried under a nitrogen gas apparatus (N-EVAP) and submitted for LC-MS analysis.

Samples were analyzed by High-Performance Liquid Chromatography and High-Resolution Mass Spectrometry and Tandem Mass Spectrometry (HPLC-MS/MS). Specifically, system consisted of a Thermo Q-Exactive in line with an electrospray source and an Ultimate3000 (Thermo) series HPLC consisting of a binary pump, degasser, and auto-sampler outfitted with a Xbridge Amide column (Waters; dimensions of 4.6 mm × 100 mm and a 3.5 µm particle size). The mobile phase A contained 95% (vol/vol) water, 5% (vol/vol) acetonitrile, 20 mM ammonium hydroxide, 20 mM ammonium acetate, pH = 9.0; B was 100% Acetonitrile. The gradient was as following: 0 min, 15% A; 2.5 min, 30% A; 7 min, 43% A; 16 min, 62% A; 16.1-18 min, 75% A; 18-25 min, 15% A with a flow rate of 400 μL/min. The capillary of the ESI source was set to 275 °C, with sheath gas at 45 arbitrary units, auxiliary gas at 5 arbitrary units and the spray voltage at 4.0 kV. In positive/negative polarity switching mode, an m/z scan range from 70 to 850 was chosen and MS1 data was collected at a resolution of 70,000. The automatic gain control (AGC) target was set at 1 × 106 and the maximum injection time was 200 ms. The top 5 precursor ions were fragmented, in a data-dependent manner, using the higher energy collisional dissociation (HCD) cell set to 30% normalized collision energy in MS2 at a resolution power of 17,500. The sample volumes of 10 μl were injected. Data acquisition and analysis were performed by Xcalibur 4.0 software and Tracefinder 2.1 software, respectively (both from Thermo Fisher Scientific).

### Fluorescent glucose uptake

Cells were incubated with 20uM of 2-NBD-Glucose Fluorescent glucose uptake probe (Abcam, ab146200) in glucose-free medium for 20min at 37°C. After washing, glucose uptake was monitored using a Laser Scanning confocal microscope (LSM780, Carl Zeiss) through a 10X and 20X objective. The mean intensity of glucose in the cell area was measured with ImageJ software (NIH) as above on z-projections, after background correction.

### Sorbitol, Lactate, Pyruvate and Glucose concentration measurement

The levels of selected metabolites were measured by commercial kits to confirm the results of metabolic profiling. These include the glucose colorimetric assay kit (BioVision), the lactate colorimetric assay kit (BioVision), the pyruvate colorimetric assay kit (BioVision) and the D-sorbitol colorimetric assay kit (BioVision) The manufacturers’ protocols were followed. Cell number was determined in concurrent experiment run in parallel, averaged per condition, and the metabolite consumption/production rates were calculated per cell. The cell numbers were measured from duplicate treatment plates to determine the proliferation rate, and the metabolite flux was determined with the following formula:

Uptake/secretion rate =Δ metabolite/(Δ time * average cell number) Average cell number = Δ cell number/(growth rate * Δ time)
Uptake/secretion rate = (Δ metabolite/Δ time) * (growth rate * Δ time/Δ cell number) = (Δ metabolite/Δ cell number) * growt rate
Growth rate [1/h] = LN(cell number T1) – LN(cell number T0)/time (T1) - time(T0)

### Atomistic simulations

The potential of mean force (PMF) was estimated by performing a set of atomistic Umbrella Sampling Molecular Dynamics (MD) simulations for the most representative pairs of amino acids contained in FUS and TAZ protein sequences. The simulations are performed with explicit water (TIP4P-D) and ions using the a99SB-*disp* force field^26^ for protein, water and ions, and using the GROMACS 2018 MD package^47^. Acetyl and N-methyl capping groups were added to the termini of each amino acid. The amino acid structure is held by imposing positional restraints on all the heavy atoms (1000 kJ mol^-1^ nm^-2^) in all directions of space (except for the dissociating amino acid in the pulling direction). As reaction coordinate, we use the center-of-mass (COM) distance between the two amino acids. Umbrella windows are spaced approximately every 0.6 Å along the reaction coordinate. To sample the steep potential, a spring constant of 6000 kJ mol^-1^ nm^-2^ is employed.

25 ns per umbrella window are simulated, to obtain uncertainties below 1 k_B_T. For the integration of the equations of motion, we used the Verlet algorithm^48^ with a time step of 1 fs. To constrain bond lengths and angles, we used the LINCS algorithm^49^ with an order of 8 and 2 iterations. The cut-off distance for the Coulomb and van der Waals interactions was chosen at a conservative value of 1.4 nm. For electrostatics, we used Particle Mesh Ewald (PME) of 4th order with a Fourier spacing of 0.12 nm and an Ewald tolerance of 1.5·10^-5^. The simulations were performed in the NpT ensemble. The temperature and pressure were kept constant using a Bussi-Donadio-Parrinello (V-rescale in GROMACS) thermostat^50^ at T = 300 K (with 1 ps relaxation time) and a Parrinello–Rahman isotropic barostat^51^ at p = 1 bar (with a 1 ps relaxation time), respectively. A NaCl concentration of 0.15 M was used throughout.

Each simulation system comprises of a box of approximately 6 x 6 x 8 nm^3^. We used approximately 7500 water molecules. All our systems were electroneutral (i.e., the total net charge was zero). After solvation of the amino acids, we perform an energy minimization with a force tolerance of 1000 kJ mol^-1^ nm^-1^, followed by a short, 1000 ps NpT equilibration both with positional restraints (9000 kJ mol^-1^ nm^-2^) for the heavy atoms of the chains in the three directions of space. The analysis of the simulations was carried out using the WHAM^52^ analysis implemented in GROMACS. The first 10% of the simulations was discarded as equilibration time.

PMF calculations are performed in aqueous solution and, additionally, in presence of sorbitol, in the concentration range 0-100mM, and glucose, between 0 and 10mM. Sorbitol and glucose topologies were obtained from ACPYPE server^53^, which utilizes the antechamber package for Amber.

### Sequence-dependent residue resolution simulations

The phase diagrams of TAZ and FUS are calculated by means of direct coexistence (DC) simulations^30^ using the molecular dynamics (MD) LAMMPS package^54^. Proteins are modeled through the CALVADOS2 force field, a sequence-dependent implicit-solvent coarse-grained model recently parameterized by Tesei *et al.*^29^. The model has a resolution of one bead per amino acid. The intrinsically disordered regions (IDRs) of the proteins are simulated as fully flexible polymers whereas the structured globular domains are modeled as rigid bodies using the conformation Protein Data Bank (PDB) crystalline structure and the Alpha Fold prediction. For FUS one globular domain is located in residues from 285–371 (PDB code: 2LCW) and the second one from 422–453 (PDB code: 86G99), whereas for TAZ we included the globular domains in residues 24-37, 130-152, and 227-264 according to the structure predicted by Alpha Fold (PDB code: Q9GZV5). Structured global domains are integrated by using the rigid body integrator of LAMMPS^54^. In addition, the hydrophobic interactions of the structured globular domains are scaled down by 30% to account for the ‘buried’ amino acids as proposed by Krainer et al.^55^. The total interaction energy per bead can be expressed as:

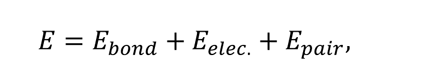

where *E*_*elec*_ and *E*_*pair*_ affect only non-bonded beads and *E*_*bond*_is only applied to bonded consectuive beads across the sequence.

Subsequent beads are linked by a harmonic potential:

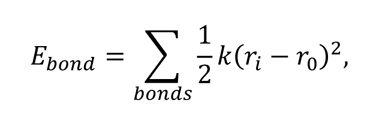

where *k* = 8033 *kk* *mol*^−1^*nm*^−2^ is the bond constant and *r*_0_ = 3.81Å is the equilibrium bond length. Electrostatic interactions are modelled via the Debye-Hückel potential to account for the salt screening:

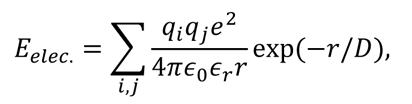

where *q*_*i*_, *q*_*j*_ account for the charges of the *i* and *j* interacting beads respectively, *e* is the elementary charge, *r* is the distance between the beads, 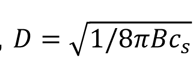 is the Debye length of an electrolyte solution of ionic strength *c*_*S*_ = 150*mM* and *B*(*ε*_*i*_) is the Bjerrum length. The electrostatic potential is truncated by a cutoff of *r*_*c*_ = 40Å. The action of the implicit solvent is implemented through the temperature-dependent dielectric constant of water as expressed by the following empirical relationship^56^:

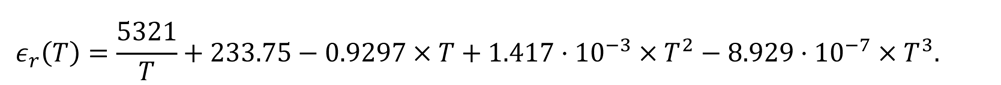

Hydrophobic interactions are parameterized via the Ashbaugh-Hatch potential^57^:

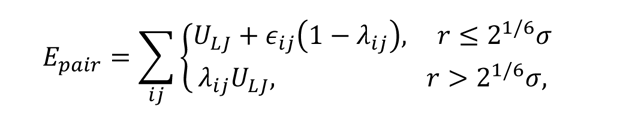

where *ε*_=*ij*_ ≡ *ε* = 0.8368*kk* *mMmol*^−1^, *λ*_*ij*_ and *λ*_*j*_ are the parameters that account for the hydrophobicity of the *i*th and *j*th interacting particles respectively, being *λ*_*ijj*_ = (*λ*_*ij*_ + *λ*_*j*_)/2. and *U*_*LJ*_ represents the standard Lennard-Jones potential:

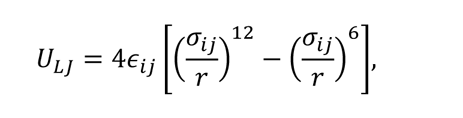

where the parameters *σ*_*ij*_ and *σ*_*ij*_ represents the excluded volume of the different residues (being *σ*_*ijj*_ = (*σ*_*ij*_ + *σ*_*j*_)/2and *r* the distance between the *ij* particles). Similar to the electrostatic interactions, the Ashbaugh-Hatch interactions are truncated at a cutoff distance of *r*_*c*_ = 40Å. The precise values of λij and σij for each pairwise interaction are reported in ^29^

For the sequence-dependent CG simulations mimicking the presence of sorbitol and glucose, we parameterize the hydrophobic interactions (e.g., λ values) according to our PMF calculations presented in **Extended Data Fig. 6**. We scale the parameter λ by the ratio between the depth of the PMF global minima in absence vs. presence of sorbitol or glucose respectively. Furthermore, this ratio is also renormalized to account for the real probability of sorbitol and glucose interacting with the amino acids inside the protein condensates. Within FUS condensates the amino acid concentration is approximately ∼1.5 M^58^ (and expectedly of the same order in TAZ droplets), which compared to our PMF calculations (where the amino acid concentration is ∼0.1 M) is roughly 15 times greater. Therefore, we rescale the PMF minima ratio between sorbitol (or glucose) compared to their absence in the solution by a factor 15.

From the PMF simulations, we reparametrized λi of glycine, alanine, glutamine, and proline, as well as the electrostatically charged interactions based on the Arg-Arg, Asp-Arg and Glu-Arg. Moreover, cation-π interactions (including Arg-Tyr, Arg-Phe and Arg-Trp) were reparametrized based on the Tyr-Arg PMF. The Tyr-Phe, Tyr-Trp and Tyr-Tyr π-π interactions were also reparametrized following the Tyr-Tyr PMF result. Once we obtain the new set of λi values at the maximum sorbitol (100 mM) and glucose (10 mM) concentrations, we linearly interpolate the parameters for intermediate concentrations using the values from the maximum sorbitol or glucose concentration and those in absence of both solutes (original model). The LAMMPS input files for all sorbitol and glucose models at the different concentrations studied here can be found in our online repository (INSERT REPOSITORY). We also note that the electrostatic interaction variation upon the addition of sorbitol or glucose in the coarse-grained model has been implemented through the effective change in λi.

The phase diagrams of TAZ and FUS are calculated both alone, in presence of glucose, and in presence of sorbitol for different concentrations using the corresponding parameters for each system (see the following repository for model parameters and input files of the CG simulations; https://doi.org/10.5281/zenodo.6979617). The number of protein replicas for each system is *Nc* = 40 and *Nc* = 48 for TAZ and FUS respectively. We employ Direct Coexistence simulations^30^ to calculate the phase diagrams in the NVT ensemble using a Langevin thermostat^59^ with a relaxation time of 50ps. All our simulations are carried out on the LAMMPS MD package (version 2^nd^ of June of 2020). Integration of the globular domains of both FUS and TAZ is carried out using the LAMMPS rigid body Nosé–Hoover thermostat^60^ in combination with the Langevin thermostat. The integration timestep is set to 10 fs. In DC simulations both the condensed and diluted coexisting phases are simulated inside an extended slab-like box with periodic boundary conditions in all directions. We first simulate 10 ns to ensure a proper equilibration of the system and then we run a longer production run of 200 ns measuring the density profile along the long axis of the simulation box, from which we obtain the bulk densities corresponding to each phase. The critical point of the system is estimated using the universal scaling law of coexistence densities near a critical point^61^, and the law of rectilinear diameters^62^:

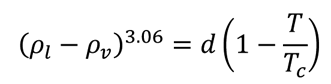

and

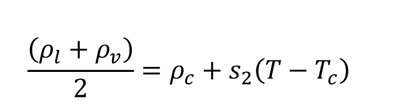

where *ρ*_*l*_ and *ρ*_*v*_ refer to the coexisting densities of the condensed and diluted phases respectively, *ρ*_*c*_ is the critical density, *T*_*c*_ is the critical temperature, and *d* and *S*_2_ are fitting parameters. The resulting critical temperatures for both proteins alone are *T*_*c*_ = 306*K* and*T*_*c*_ = 312*K* for FUS and TAZ respectively.

### FRAP

FRAP experiment were performed on Laser Scanning Confocal Microscope (LSM780 Carl Zeiss) equipped with a heated stage maintained at 37°C and the whole power of its 30mW 488 nm Argon laser through a 63X/1.4 NA oil immersion objective. Fluorescence recovery in bleached region of interest (condensate) was analyzed using the FRAP module of ZEN software (Carl Zeiss). Mobile fraction (in %) and half-time of recovery (in seconds) values were extracted for each experiment.

### Immunofluorescence

Cells were fixed with PBS/PFA 4% for 10 min and permeabilized with PBS/Triton 100X 0.2% for 5 min. After blocking with PBS/Triton/BSA 0.2% for 1h, the cells were then incubated with primary antibodies (FUS, TAZ, Gal3 or GFP; 1/100) at room temperature for 1 h. Secondary antibodies coupled with Alexa-594 and/or Alexa-488 (A-11012, A-11001, Life technologies) were used at 1/500. Nuclei were counterstained with DAPI (Sigma-Aldrich). Images were obtained using a Laser Scanning confocal microscope (LSM780, Carl Zeiss) through a 63X/1.4 NA oil immersion objective.

### Circularity measurement

The circularity index [4p(area/perimeter^2^)] of condensate shape was quantified as previously described^63^. For application of circularity measurements of condensates, a freehand selection option in ImageJ software was used to delineate cell outline. Circularity index values were assigned to cell outlines.

### Condensate intensity and number

To quantify condensate intensity and number we selected condensate with a size > 100nm and excluded all the condensates with a circularity index <0.8. The mean intensity of condensate (Alexa-488 or Alexa-594 fluorescence) in the cell area was measured with ImageJ software (NIH) on z-projections, after background correction. Number of condensates in the cell or per cell was calculated using the Fiji Analyze Particles plugin as above on z-projections after threshold.

### ChIP-qPCR

MDA-MB-231 cells were cultivated on indicated substrate for 48h. Cells were dual cross-linked with 2 mM disuccinimidylglutarate (DSG) for 45 minutes and then in 1% paraformaldehyde for 15 minutes at room temperature. Fixed cells were lysed in 10 ml of Lysis Buffer 1 [50 mM HEPES (pH 7.5), 140 mM NaCl, 1 mM EDTA, 0.1% IGEPAL 630 (Sigma Aldrich)], containing 0.05% Triton X100, 2.5 % glycerol and supplemented with 1X protease inhibitor cocktail (Roche) for 10 minutes on ice, followed by incubation in Buffer 2 [0.1 M Tris HCl (pH 8) and 200 mM NaCl with protease inhibitors] for 15 minutes at room temperature. Chromatin was sonicated at 30% of amplitude for 10 minutes (10 cycles of 1 minute). The samples were centrifuged (2X 14,000 g for 5 minutes each), and soluble chromatin was transferred to a fresh tube. Crosslinked DNA after sonication was precipitated with 5μg of anti-TAZ antibody (Cell Signaling, Rabbit mAb #70148) or non-immune rabbit IgG (ab27472, Abcam) overnight at 4°C. Chromatin/antibody complex was pulled down with PureProteome™ Protein G Magnetic Beads (Millipore) and washed in the low-and high-salt buffers. After crosslinking reversion (65°C for 4 hours) and Proteinase K treatment, chromatin was purified by phenol-chloroform extraction and ethanol precipitation. Precipitated DNA was analyzed by qPCR using primers generated for predicted TEAD binding sites.

### Inhibition of Lox in breast cancer mouse model

All animal husbandry, procedures performed, and assay endpoints were in accordance with protocols approved by the local committee of the host institute and by the Institutional Animal Care and Use Committee (CIEPAL AZUR committee,) University Cote d’Azur, Nice, France. 8 weeks old female BALB/c mice were obtained from Janvier Laboratory. Metastatic mouse (BALB/c) 4T1 breast cancer cells were implanted into the right fourth mammary fat pad in 10 μl Matrigel of 8-week-old female BALB/c mice. After 9 days, mice with palpable tumor (5-10mm3) were randomly assigned to treatment groups. β-aminopropionitrile (BAPN; 100 mg/kg/d; Sigma-Aldrich) was administered in drinking water. Mice were randomly assigned to experimental groups.

### Human studies

Tissue microarray containing BC081120e were provided by US Biomax, which contained a total of 100 cancer cases of breast cancer and only a single punch for the one patient. Inclusion criteria included female sex, original histological diagnosis of invasive breast carcinoma, validation of the status of ER, PR, and HER2 as well as availability of clinical pathological data. Experiments with tumor tissues were conducted in compliance with the Helsinki Declaration and approved by the Ethics Committee of our University.

### Immunohistochemistry and immunofluorescence of mammary tumor sections

Mammary tumor sections (5μm) were deparaffinized and high-temperature antigen retrieval was performed, followed by blocking in TBS/BSA 5%, 10% donkey serum and exposure to primary antibody and biotinylated secondary antibody (Vectastain ABC kit, Vector Labs) for immunohistochemistry or Alexa 488, 568 and 647-conjugated secondary antibodies (Thermo Fisher Scientific) for immunofluorescence. Primary antibodies against, Ki67 (ab16667; 1/100) and α-SMA (ab32575; 1/1000 or ab21027; 1/300) were purchased from Abcam. A primary antibody against TAZ (sc-48805, 1/100) was purchased from Santa Cruz Biotechnology. A primary antibody against FUS (ab23439; 1/100), was purchased from Abcam. A primary antibody against Galectin-3 (60207-1-Ig; 1/100), was purchased from Proteintech. Pictures were obtained using an Olympus Bx51 microscope or ZEISS LSM Exciter confocal microscope. Intensity of staining was quantified using ImageJ software (NIH) as previously described (Bertero et al., 2019a). All measurements were performed blinded to condition.

### Picrosirius red stain and quantification

Picrosirius red stain was achieved through the use of 5 mm paraffin sections stained with 0.1% Picrosirius red (Direct Red80, Sigma-Aldrich) and counterstained with Weigert’s hematoxylin to reveal fibrillar collagen. The sections were then imaged and quantified as previously described^38^. Two authors, blinded to each other’s assessment, scored the slides using the Quick Score method (as described^64^:) to determine the ECM remodeling status (Picrosirius red staining) within the tumor.

### Immunohistochemical staining and quantification methods of human samples

After deparaffinization and rehydration, microwave antigen retrieval was performed in Na-citrate buffer (10mM, pH6; 5min at 900W, 10min at 150W and 30 min at room temperature). Sections were washed three times in PBS (5min per wash). After incubation in blocking buffer for one hours (PBS 3%BSA; 10% serum; 0.3% Triton X-100), sections were incubated with primary antibody for α−SMA (ab21027; 1/300), TAZ (sc-48805, 1/100), FUS (ab23439; 1/100), Galectin-3 (60207-1-Ig; 1/100), and Ki67 (ab16667; 1/100) staining diluted in blocking buffer overnight at 4°C. After three washes in PBS/0,1% Tween 20, sections were incubated with secondary antibody diluted 1:400 in blocking buffer for 1 hour at room temperature and washed 3 times in PBS/0,1% Tween 20. Nuclei were then stained with DAPI and mounted in Permafluor (Thermo Scientific).

### Quantification and statistical analysis

Cell culture experiments were performed at least three times and at least in triplicate for each replicate. The number of animals in each group was calculated to measure at least a 20% difference between the means of experimental and control groups with a power of 80% and standard deviation of 10%. The number of unique patient samples for this study was determined primarily by clinical availability. In situ expression/histological analyses of both mouse and human tissue were performed in a blinded fashion. Immunoblot images are representative of experiments that have been repeated at least three times. Micrographs are representative of experiments in each relevant cohort. All analyses were performed using Prism 6.0 software (GraphPad, Paired samples were compared by a 2-tailed Student’s t test for normally distributed data, while Mann-Whitney U non-parametric testing was used for non-normally distributed data. For comparing more than two conditions, one-way ANOVA was used with: Bonferroni’s multiple comparison test or Dunnett’s multiple comparison test (if comparing all conditions to the control condition). A p value less than 0.05 was considered significant and marked on the graphs by asterisks. Error bars denote SEM. Correlation analyses were performed by Pearson correlation coefficient calculation. No samples/animals/patients were excluded.

## Data availability

All the data are available upon reasonable request.

## Acknowledgments

We thank the B. Mari team for advice and discussions. The author acknowledges the “Microscopie Imagerie Côte d’Azur” (MICA), GIS-IBISA multi-sites platform and particularly it’s IPMC, C3M and IRCAN (Molecular and Cellular Imaging facility PICMI) partners. **Funding:** This platform is supported by the GIS IBiSA, Conseil Départemental 06, Région PACA ARC, Cancerôpole PACA,)”. This work was supported by; the French National Research Agency ANR-18-CE14-0025, ANR-20-CE14-0006 and ANR-21-CE44-0036, the French National Cancer Institute INCA-PLBIO 21-094 and the foundation ARC pour la recherché sur le cancer PJA20191209291 (to T.B.); and, the foundation ARC pour la recherché sur le cancer PJA2021050003611 and Cancerôpole PACA, Emergence grant 2021 (to S.T.), and grant from the National Institutes of Health R01GM143334 (I.B-.S.). I.S.-B. acknowledges funding from the Derek Brewer scholarship of Emmanuel College and EPSRC Doctoral Training Programme studentship, number EP/T517847/1. A.R.T. and J.R.E. acknowledge funding from the Ramon y Cajal fellowship (RYC2021-030937-I). J.R.E. aknowledges funding from the Spanish scientific plan and committee for research reference PID2022-136919NA-C33. A.R.T. and R.C.-G. acknowledge funding from the European Research Council (ERC) under the European Union Horizon 2020 research and innovation programme (grant agreement 803326). Atomistic and coarse-grained simulations were performed using resources provided by the Cambridge Tier-2 system operated by the University of Cambridge Research Computing Service (http://www.hpc.cam.ac.uk) funded by EPSRC Tier2 capital grants EP/P020259/1-CS092 and EP/P020259/Su111.

## Author contributions

ST and TB conceived and designed the experiments. ST, NR, CC, SB, SC and TB performed the majority of the experiments. WMO, IB-S, and TB, provided the experimental infrastructure. FB and SA provide critical insight for microscopy experiments. WMO performed and analysed steady state metabolomics experiments. IB-S and BPO performed and analysed ^13^C6-glucose tracing experiments. ART, IS-B, RC-G and JRE performed and analysed atomistic simulations. ST and TB wrote the manuscript. All authors participated in interpreting the results and revising the manuscript.

## Additional information

**Extended Data figure 1:** Matrix stiffening promotes FUS biomolecular condensates.

**Extended Data figure 2:** Matrix stiffening promotes Galectin 3 biomolecular condensates.

**Extended Data figure 3:** Matrix stiffening promotes endogenous biomolecular condensates.

**Extended Data figure 4:** Matrix stiffening promotes biomolecular condensates in primary endothelial cells.

**Extended Data figure 5:** Matrix stiffening rewire glucose to the polyol pathway.

**Extended Data figure 6:** Atomistic simulations

**Extended Data figure 7:** Sequence-dependent residue resolution simulations

**Extended Data figure 8:** Physiological sorbitol concentrations force biomolecular condensates.

**Extended Data figure 9:** Intracellular sorbitol concentration regulates protein phase behavior.

**Extended Data figure 10:** Matrix stiffening forces biomolecular condensates in breast tumor

