## Extended Data for "Mechano-dependent sorbitol accumulation supports biomolecular condensate"

**Extended Data figure 1: Matrix stiffening promotes FUS biomolecular condensates.**

**(a-d)** MDA-MB-231 cells plated on the indicated substrate and treated with 10µM Y-27632 (Y27) or 5µM PF-573228 (PF) for 1hour **(d)**. **(a,d)** Representative immunofluorescence images and quantification of intensity, number per cell of GFP-TAZ condensates and number of cell with condensates. Scale bar=10 µm. **(b)** Representative images and FRAP curves (left) and quantification of diffusion rate (t ½) and mobile fraction (right) of GFP-TAZ**.** White arrows indicate condensates. **(c)** Circularity quantification of TAZ protein condensates. In all the panels n>30 cells from 3 independent experiments were analyzed. ns=not significant; *P<0.05; **P<0.01; ***P<0.001; ****P<0.0001; **(a, d)** Bonferroni's multiple comparison test; data are mean ± s.e.m.

**Extended Data figure 2: Matrix stiffening promotes Galectin 3 biomolecular condensates.**

**(a-d)** MDA-MB-231 cells plated on the indicated substrate and treated with 10µM Y-27632 (Y27) or 5µM PF-573228 (PF) for 1hour **(d)**. **(a,d)** Representative immunofluorescence images and quantification of intensity, number per cell of GFP-Gal3 condensates and number of cell with condensates. Scale bar=10 µm. **(b)** Representative images and FRAP curves (left) and quantification of diffusion rate (t ½) and mobile fraction (right) of GFP-Gal3**.** White arrows indicate condensates. **(c)** Circularity quantification of Gal3 protein condensates. In all the panels n>30 cells from 3 independent experiments were analyzed. ns=not significant; *P<0.05; **P<0.01; ***P<0.001; ****P<0.0001; **(a, d)** Bonferroni's multiple comparison test; data are mean ± s.e.m.

**Extended Data figure 3: Matrix stiffening promotes endogenous biomolecular condensates.**

**(a-c)** MDA-MB-231 cells plated on the indicated substrate and treated with 5% of 1,6-Hexanediol (1,6 HD) for 5min. Representative immunofluorescence images and quantification of intensity, number per cell of endogenous TAZ **(a)**, FUS **(b)** or Gal3 **(c)** condensates and number of cell with condensates. White arrows indicate condensates. Scale bar=10 µm. In all the panels n>30 cells from 3 independent experiments were analyzed. ns=not significant; *P<0.05; **P<0.01; ***P<0.001; ****P<0.0001; **(a-f)** Bonferroni's multiple comparison test; data are mean ± s.e.m.

**Extended Data figure 4: Matrix stiffening promotes biomolecular condensates in primary endothelial cells.**

**(a-f)** PAEC cells plated on the indicated substrate and treated with 10µM Y-27632 (Y27) or 5µM PF-573228 (PF) for 1hour **(b,d,f)**. **(a-f)** Representative immunofluorescence images and quantification of intensity, number per cell of GFP-FUS **(a-b)**, GFP-TAZ **(c-d)** or GFP-Gal3 **(e-f)** condensates and number of cell with condensates. White arrows indicate condensates. Scale bar=10 µm. In all the panels n>30 cells from 3 independent experiments were analyzed. ns=not significant; *P<0.05; **P<0.01; ***P<0.001; ****P<0.0001; **(a-f)** Bonferroni's multiple comparison test; data are mean ± s.e.m.

**Extended Data figure 5: Matrix stiffening rewire glucose to the polyol pathway.**
**(a-m)** MDA-MB-231 cells plated on the indicated substrate and treated with 10µM Y-27632 (Y27) or 5µM PF-573228 (PF) for 1hour. **(a-b)** Heatmap **(a)** and pathway enrichment analysis **(b)** of significantly (FDR<1%; P<0.05) modulated intracellular metabolites. Red: metabolite from polyol pathway. **(c)** Lactate and Pyruvate ratio in cells. **(d)** Quantification of lactate secretion. **(e)** Quantification of glucose uptake. **(f)** Quantification of GLUT1 mRNA level. **(g)** Quantification of intracellular glucose**. (h-m)** Cells were exposed for 1 hour or 24 hours to $^{13}C_6$-glucose. $^{13}C_6$-glucose incorporation in intracellular Fructose-/Glucose- 6 Phosphate **(h),** Sorbitol **(i),** Dihydroxyacetone phosphate **(j),** Fructose 1,6 bis Phosphate (**k**), Pyruvate (**l**), and lactate **(m).** ns=not significant; *P<0.05; **P<0.01; ***P<0.001; ****P<0.0001; **(c-m)** Bonferroni's multiple comparison test; data are mean ± s.e.m.

**Extended Data figure 6: Atomistic simulations**
**(a-k)** Atomistic Potential of Mean Force (PMF) dissociation curve of the different amino acid pairs studied as a function of the centre of mass distance (COM) using the a99SB-disp force field. PMF simulations are conducted at room conditions and physiological NaCl concentration (150 mM). Black curves represent the amino acid PMF curves in absence of both glucose and sorbitol, while those in red and green are performed in presence of sorbitol (100 mM) and glucose (10 mM), respectively.

**Extended Data figure 7: Sequence-dependent residue resolution simulations**
**(a)** Phase diagram in the temperature-density plane for FUS using a sequence-dependent residue-resolution model (black), as well as for the reparameterizations performed in this work (based on the all-atom PMF calculations reported in the Extended Data Figure 6) including sorbitol at 100 mM concentration (red) and glucose at 10 mM (green). Temperatures have been renormalized by the critical temperature of FUS in absence of both solutes which is 306 K. **(b)** Variation of the critical temperature of FUS LLPS in presence of different concentrations of glucose and sorbitol as indicated in the figure.

**Extended Data figure 8: Physiological sorbitol concentrations force biomolecular condensates.**

**(a)** SDS-PAGE analysis of GFP-FUS, GFP-Gal3, GFP-TAU, GFP-SHP2, GFP-NPM-1, GFP-αSyn purified protein. **(b-g)** Representative images and quantification of intensity and number of GFP-FUS **(b),** GFP-Gal3 **(c),** GFP-TAU **(d)**, GFP-SHP2 **(e)**, GFP-NPM-1 **(f)** or GFP-αSyn **(g)** condensates with different sorbitol concentration. Scale bar=10 µm. ns=not significant; *P<0.05; **P<0.01; ***P<0.001; ****P<0.0001; **(b-g)** Bonferroni's multiple comparison test; data are mean ± s.e.m of at least n=3 independent experiments.

**Extended Data figure 9: Intracellular sorbitol concentration regulates protein phase behavior.**

**(a-f)** MDA-MB-231 **(a-c)** or PAEC **(d-f)** cells plated on the indicated substrate and treated with 10µM EBPC. Representative immunofluorescence images and quantification of intensity, number per cell of endogenous GFP-TAZ **(a-d)**, GFP-FUS **(b-e)** or GFP-Gal3 **(c-f)** condensates and number of cells with condensates. White arrows indicate condensates. Scale bar=10 µm. In all the panels n>30 cells from 3 independent experiments were analyzed. ns=not significant; *P<0.05; **P<0.01; ***P<0.001; ****P<0.0001; **(a,f)** two tailed t-test; data are mean ± s.e.m of at least n=3 independent experiments.

**Extended Data figure 10: Matrix stiffening forces biomolecular condensates in breast tumor**

**(a-b)** Representative images of primary mammary tumors are shown as isolated from a cohort of 100 patients. Representative IHC images show FUS **(a),** Gal3 **(b)** and α-SMA staining. Scale bar=100 µm. Correlations between FUS or Gal3 staining, and ECM remodeling are shown. Pearson correlation coefficients ($R^2$) are indicated. **(c-j)** Following cancer cell implantation, mice were treated with daily BAPN (100 mg/kg; n = 10) or vehicle (4T1 cells, n = 10). **(d)** Representative picrosirius red images and quantification showing the collagen remodeling. **(e)** Quantification of intratumoral sorbitol concentration. **(f,i-j)** Representative IHC images and quantification of cell with TAZ **(f)**, FUS **(i)** or Gal3 **(j)** condensates and TAZ **(f)**, FUS **(i)** or Gal3 **(j)** condensates number. **(g-h)** Correlations between TAZ staining and ECM remodeling **(g)** and between TAZ staining and proliferation **(h)** are shown. Pearson correlation coefficients ($R^2$) are indicated. Scale bar=100 µm. ***P<0.001; **(e-f, i-j)** two tailed t-test; data are mean ± s.e.m of at least n=3 independent experiments.
