## Extended Data Figures for "Mechano-dependent sorbitol accumulation supports biomolecular condensate"

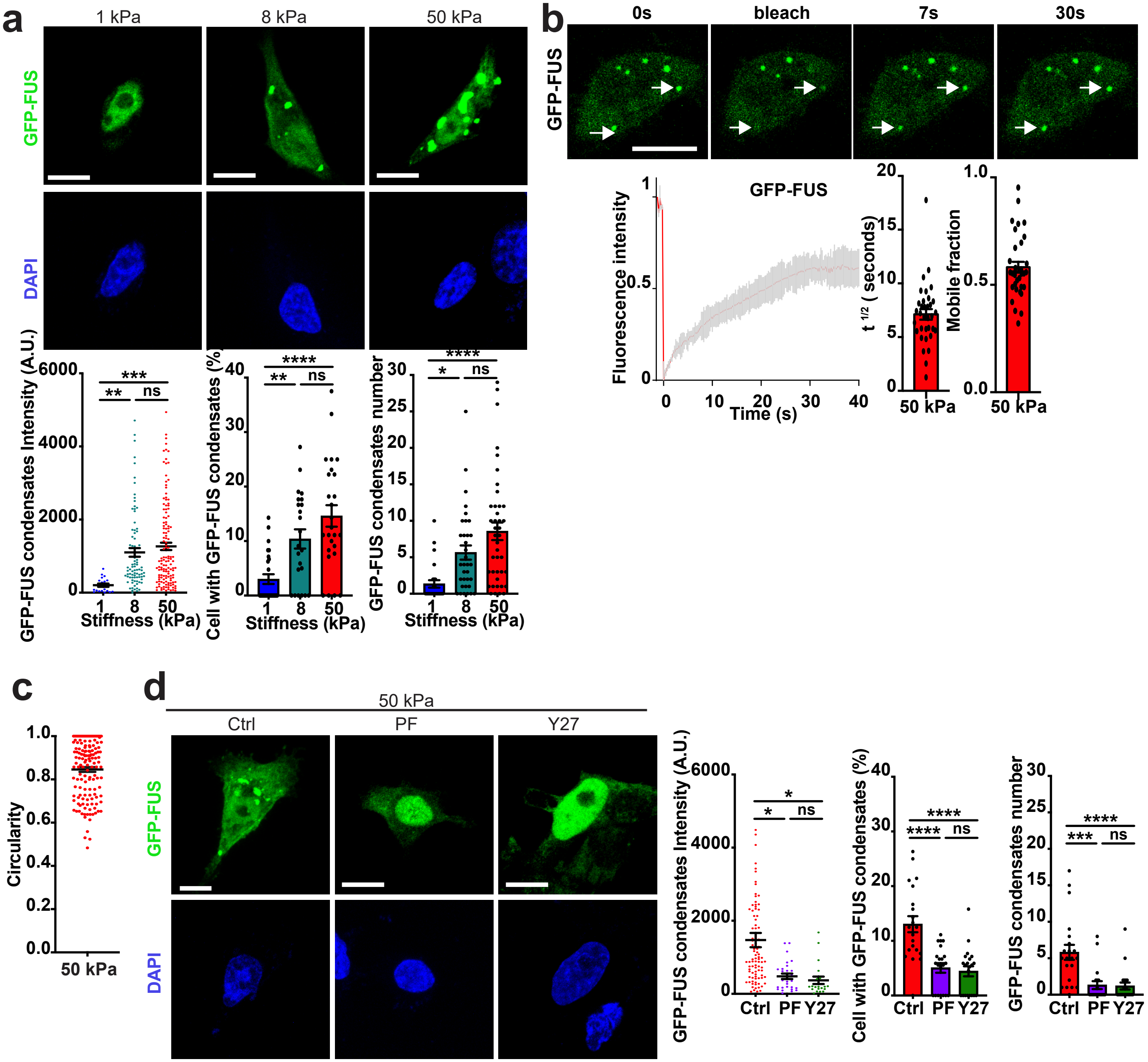


**Extended Data figure1**

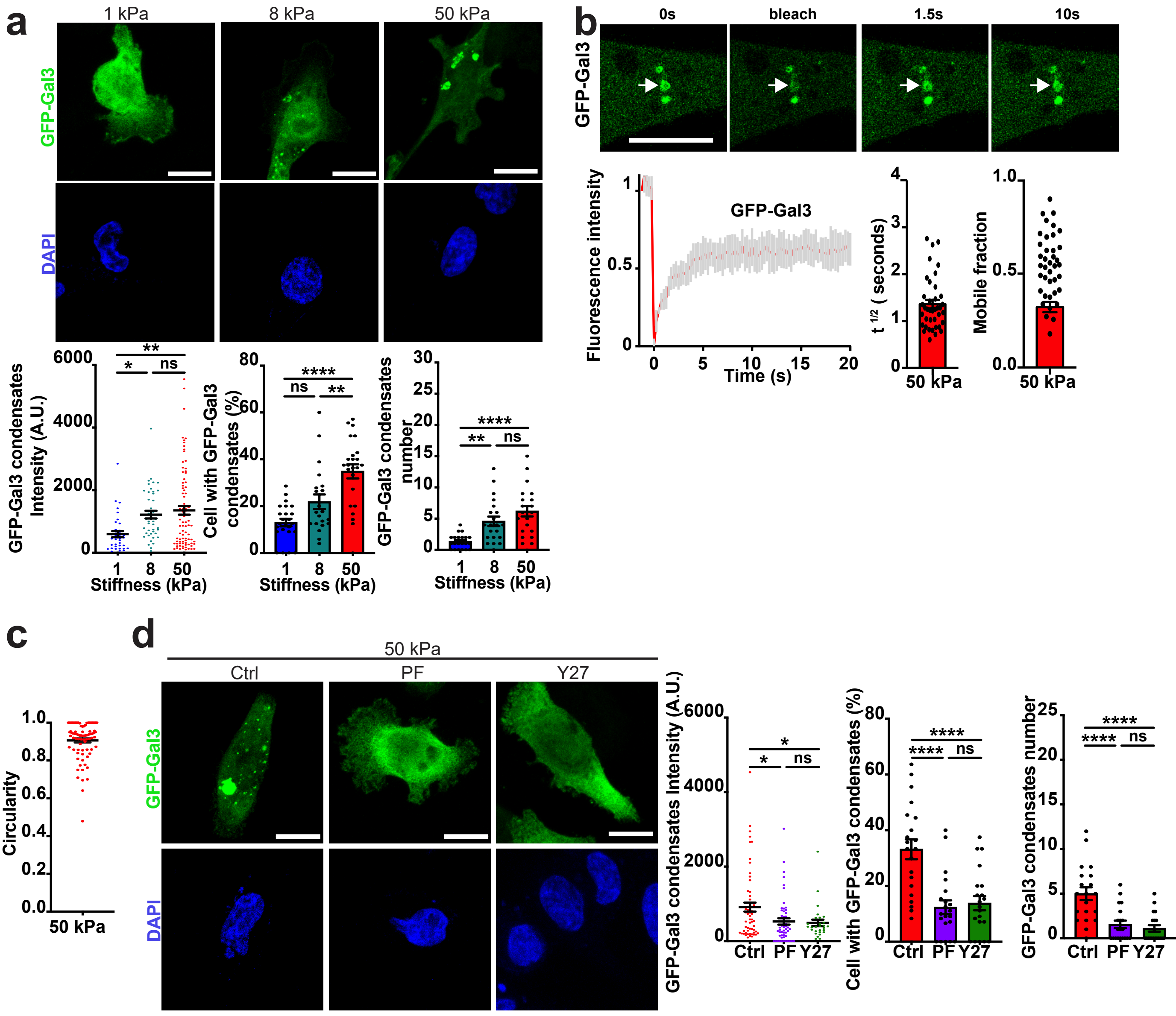


Extended Data figure 2

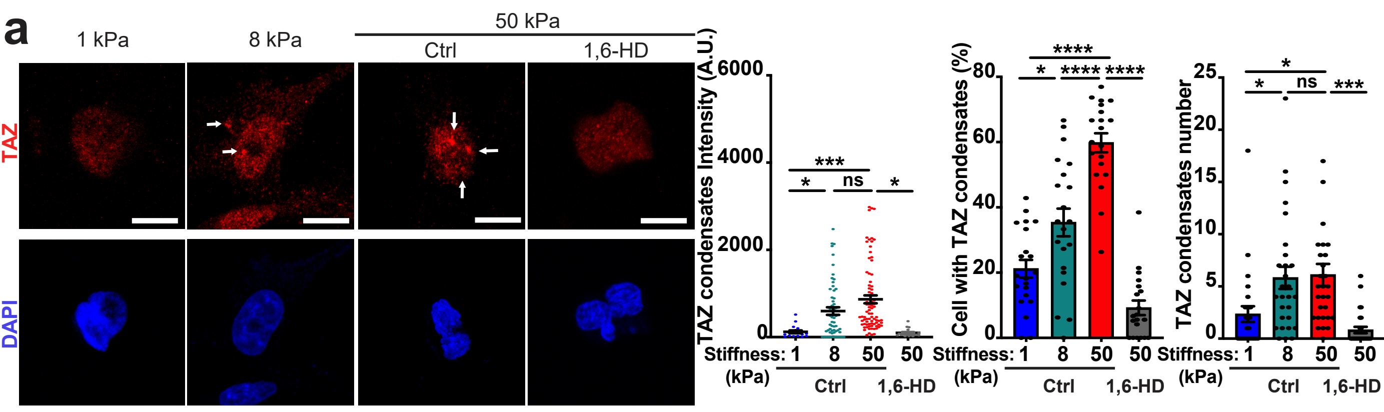


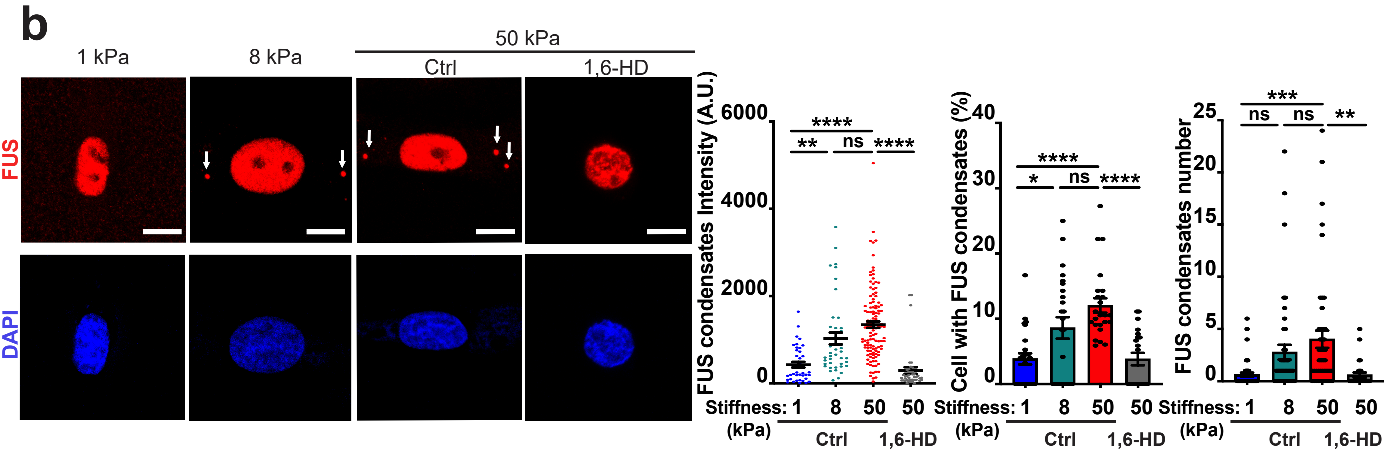


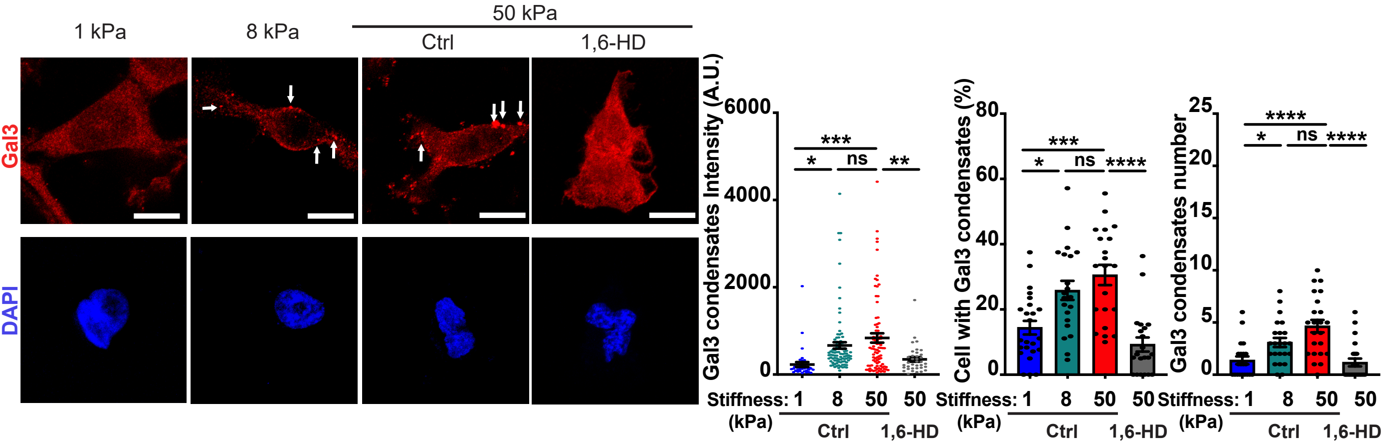


**Extended Data figure 3**

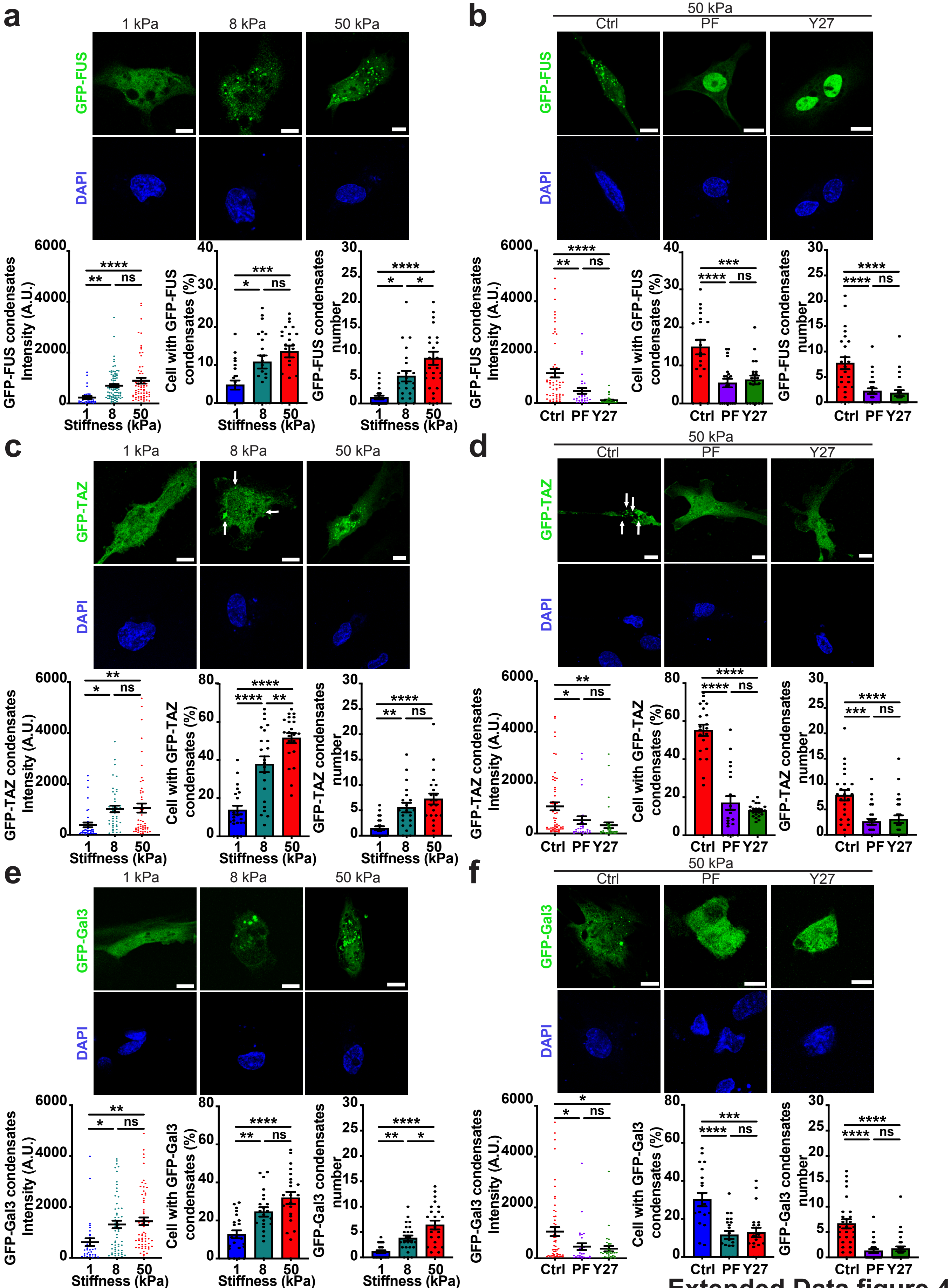


Extended Data figure 4

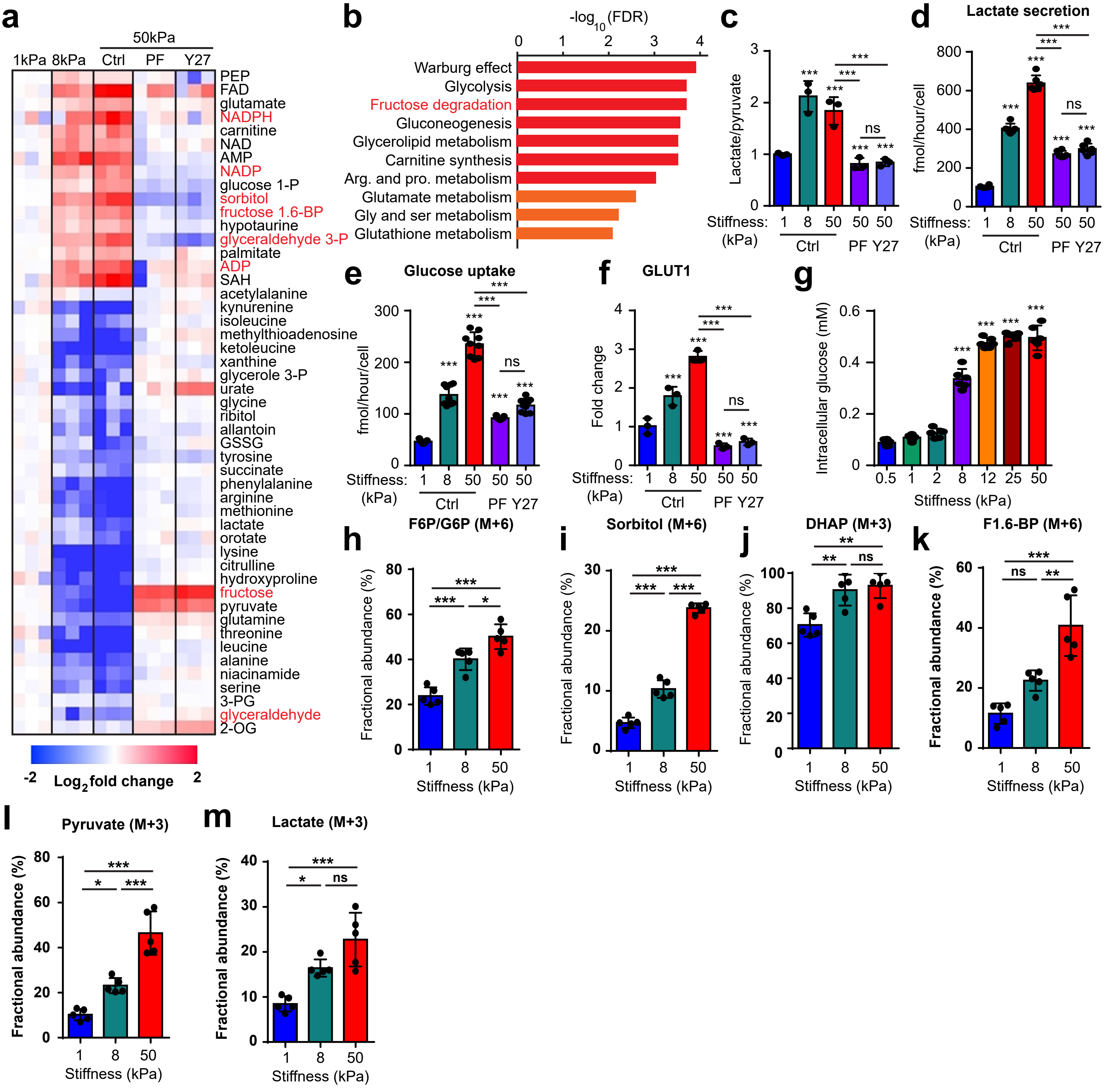


**Extended Data figure 5**

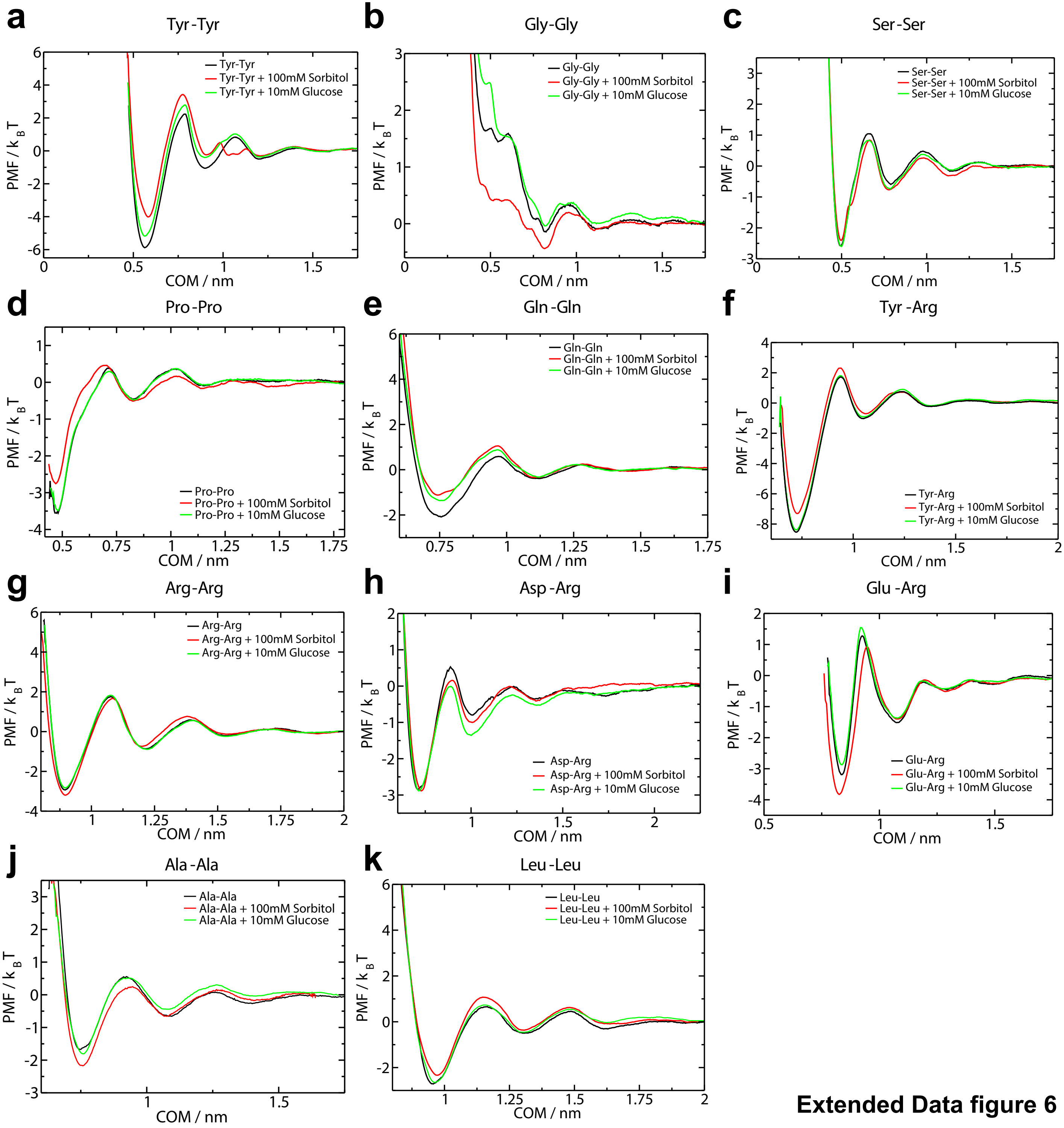


**Extended Data figure 6**

**a**

**b**

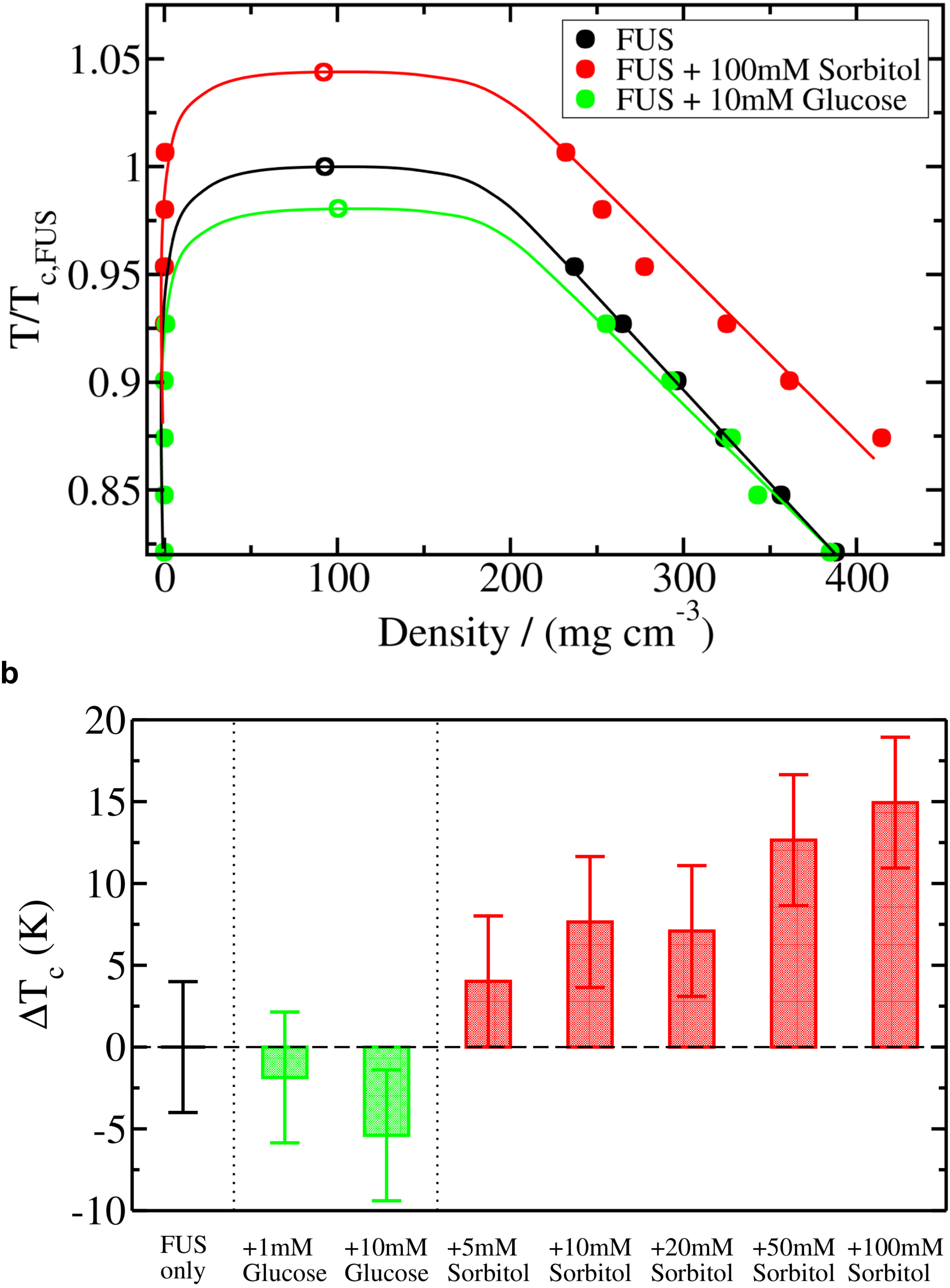


**Extendad Data figure 7**

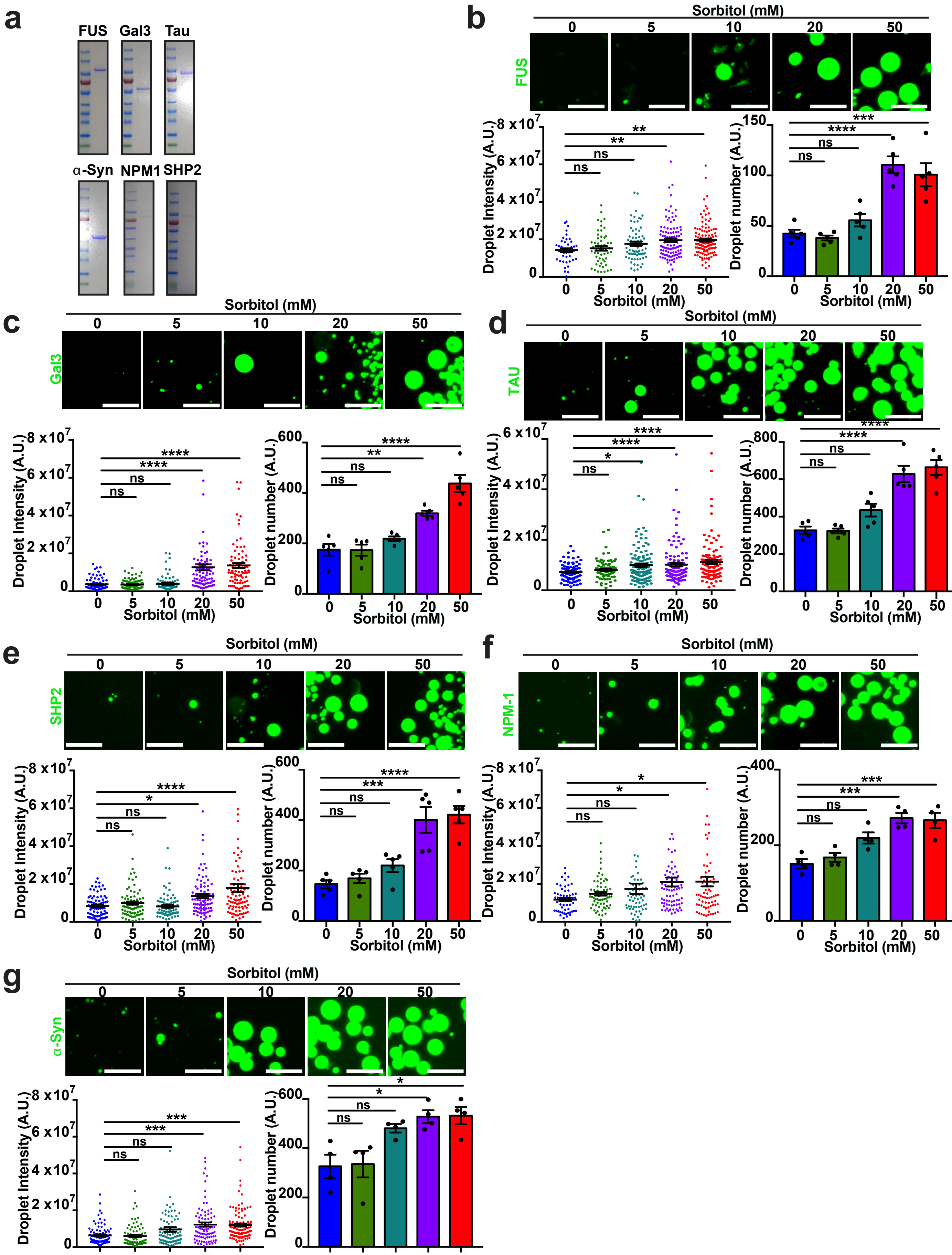


Extended Data figure 8

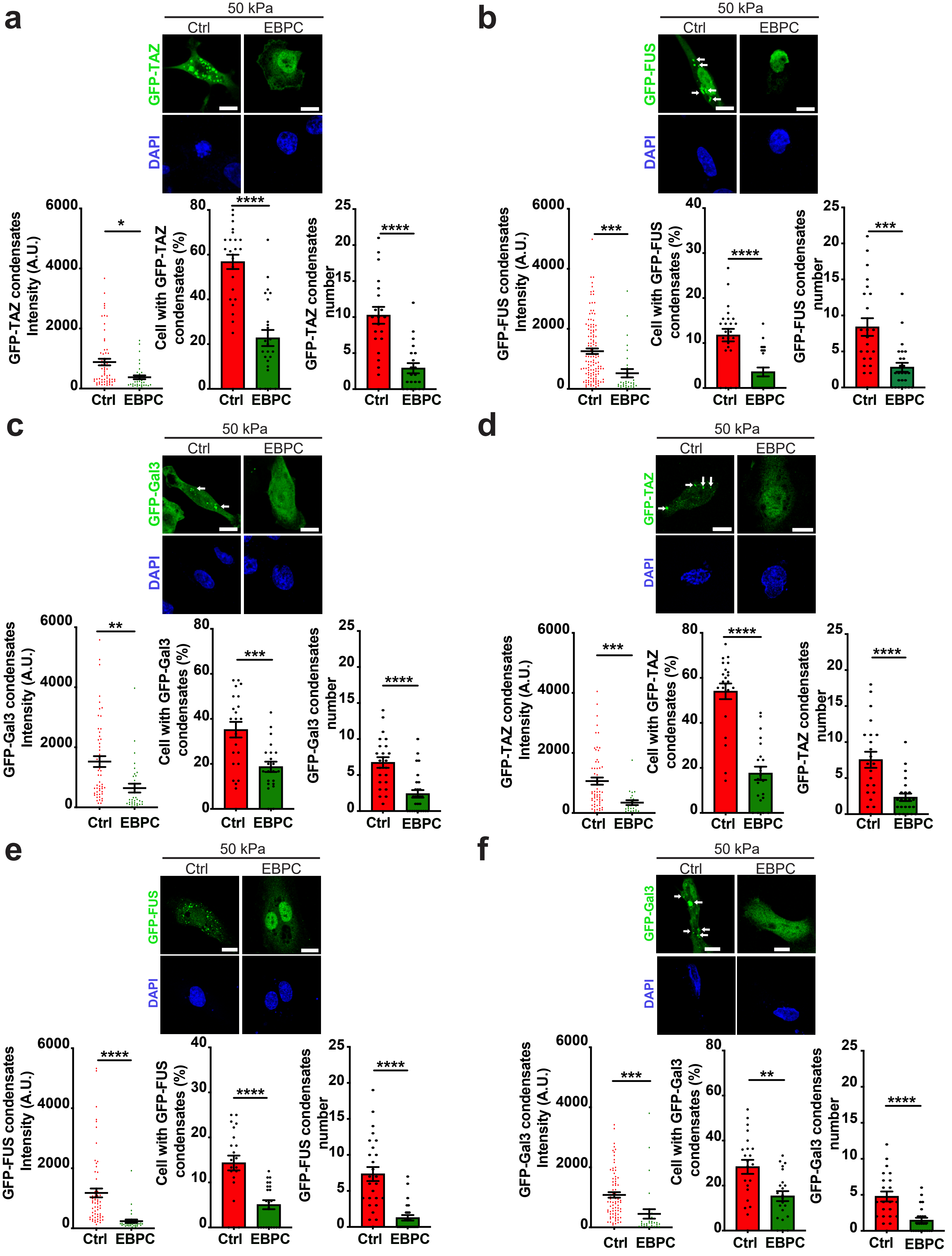


Extended Data figure 9

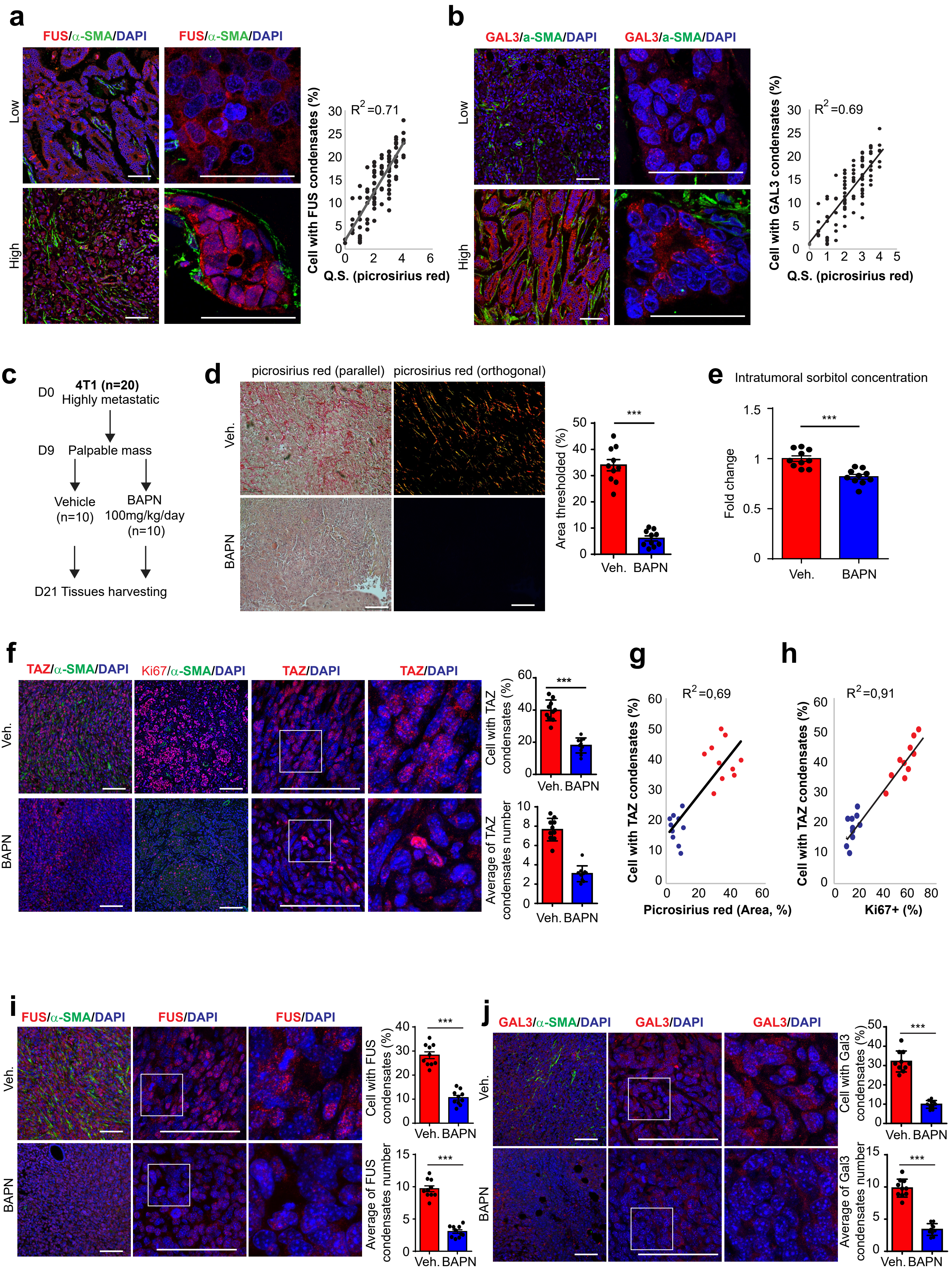
a
FUS/α-SMA/DAPI
FUS/α-SMA/DAPI
Low
High
$R^2$=0.71
Cell with FUS condensates (%)
Q.S. (picrosirius red)
b
GAL3/a-SMA/DAPI
GAL3/a-SMA/DAPI
Low
High
$R^2$=0.69
Cell with GAL3 condensates (%)
Q.S. (picrosirius red)
c
D0 4T1 (n=20) Highly metastatic
D9 Palpable mass
Vehicle (n=10)
BAPN 100mg/kg/day (n=10)
D21 Tissues harvesting
d
picrosirius red (parallel)
picrosirius red (orthogonal)
Veh.
BAPN
Area thresholded (%)
***
e
Intratumoral sorbitol concentration
Fold change
***
Veh.
BAPN
f
TAZ/α-SMA/DAPI
Ki67/α-SMA/DAPI
TAZ/DAPI
TAZ/DAPI
Veh.
BAPN
Cell with TAZ condensates (%)
Average of TAZ condensates number
***
g
$R^2$=0,69
Cell with TAZ condensates (%)
Picrosirius red (Area, %)
h
$R^2$=0,91
Cell with TAZ condensates (%)
Ki67+ (%)
i
FUS/α-SMA/DAPI
FUS/DAPI
FUS/DAPI
Veh.
BAPN
Cell with FUS condensates (%)
Average of FUS condensates number
***
j
GAL3/α-SMA/DAPI
GAL3/DAPI
GAL3/DAPI
Veh.
BAPN
Cell with Gal3 condensates (%)
Average of Gal3 condensates number
***
